# Thermodynamic Forces from Protein and Water Govern Condensate Formation of an Intrinsically Disordered Protein Domain

**DOI:** 10.1101/2023.09.05.556343

**Authors:** Saumyak Mukherjee, Lars V. Schäfer

## Abstract

Liquid-liquid phase separation (LLPS) can drive a multitude of cellular processes by compartmentalizing biological cells via the formation of dense liquid biomolecular condensates, which can function as membraneless organelles. Despite its importance, the molecular-level understanding of the underlying thermodynamics of this process remains incomplete. In this study, we use atomistic molecular dynamics simulations of the low complexity domain (LCD) of human fused in sarcoma (FUS) protein to investigate the contributions of water and protein molecules to the free energy changes that govern LLPS. Both protein and water components are found to have comparably sizeable thermodynamic contributions to the formation of FUS condensates. Moreover, we quantify the counteracting effects of water molecules that are released into the bulk upon condensate formation and the waters retained within the protein droplets. Among the various factors considered, solvation entropy and protein interaction enthalpy are identified as the most important contributions, while solvation enthalpy and protein entropy changes are smaller. These insights provide detailed molecular insights on the intricate thermodynamic interplay between protein- and solvation-related forces underlying the formation of biomolecular condensates.

## Introduction

The interior of a biological cell is densely packed with biomolecules,^1,2^ with fractions ranging up to 30% in volume or concentrations of 300 mg mL^−1^. In this intricately crowded environment, a multitude of biomolecules interact with each other.^1,3–5^ Biological functions can be modulated or even governed by such interactions.^6–8^ For example, protein–protein interactions are key for maintaining cellular homeostasis,^9^ can modulate protein stability,^2^ and may sometimes also lead to aggregation, with associated complications and diseases such as cataracts, Alzheimer’s, Parkinson’s, frontotemporal dementia and amyotrophic lateral sclerosis.^10–16^ Hence, understanding the nature of these interactions and unraveling the molecular driving forces that are at play in such crowded biomolecular environments is of fundamental importance, as it might form the basis for targeted manipulation.^17^

Apart from the biomolecules themselves, their interactions with and coupling to water, another major component of the cell, are also crucial.^18–22^ Water is essential for many biomolecular processes, including aggregation/association, and the thermodynamic contributions linked to water can be substantial.^12,21,23,24^ A prime example of biomolecular association is liquid–liquid phase separation (LLPS) of proteins and nucleic acids.^25–28^ LLPS leads to compartmentalization within biological cells via formation of dense biomolecular condensates or droplets,^29,30^ which are dispersed in a more dilute environment. This process depends on a multitude of factors such as temperature, pressure, pH, cosolvents, salt concentration, etc.^28,31^

Membraneless organelles are intensely researched because they have been shown to serve as selective “microreactors” within cells in which specific biochemical reactions can take place. These can be pivotal for a plethora of cellular processes, such as RNA splicing, receptor-mediated signalling, and mitosis.^32–36^ The dynamic nature and liquid-like character of these condensates allow for the efficient exchange of components with the surroundings.^37^

Several proteins have been shown to undergo LLPS.^38,39^ The fused in sarcoma (FUS) RNA-binding protein is one such protein that was found to form intracellular condensates.^40–42^ FUS is crucial for RNA shearing and transport, DNA repair, micro-RNA processing, gene transcription and regulation.^43–47^ Human FUS is a 525-residue protein with an N-terminal intrinsically disordered low complexity domain (LCD). This LCD region is responsible for the LLPS of FUS,^48–50^ primarily mediated by multivalent interactions.

Solvation effects are important for protein condensate formation.^23^ The present work is based on the consideration that when proteins come close to each other in the condensate, some of the water molecules in the vicinity of the protein surfaces are replaced by other protein moieties. Hence, these water molecules are released into the surrounding dilute phase, resulting in a partial dewetting of the protein surfaces. It was speculated that this water release could be associated with an entropy gain.^23^ Indeed, experimental phase transition data for the N-terminal part of Ddx4^51^ and for tau-RNA droplet formation,^52^ analyzed within the framework of the Flory-Huggins model of polymer phase transitions, suggests that the phase separation entropy was favorable. Recent THz spectroscopy experiments were interpreted along the same lines.^53,54^ However, at the same time, the condensates also retain a substantial amount of water,^49,55^ which could experience entropy loss due to increased confinement. Figure 1 schematically illustrates the idea of the interplay, or “tug-of-war”, of released and retained water, which also forms the basis of the present work.

**Figure 1:**
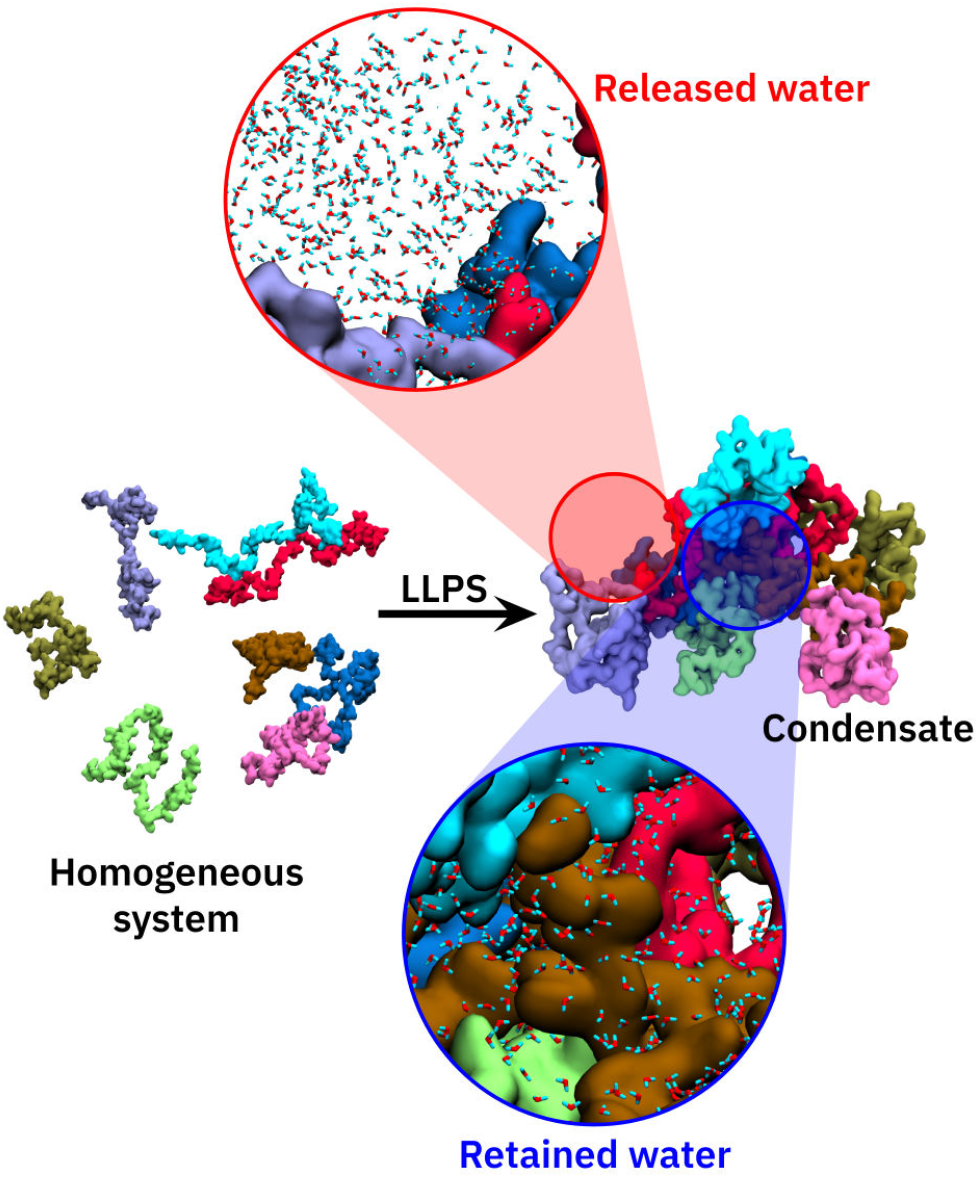
Illustration of the process of protein condensate formation via liquid–liquid phase separation. The zoomed-in views highlight the water molecules that are released into a bulk-like environment (top) outside the protein condensate and the ones that are retained inside the condensate (bottom).

Due to the challenges associated with studying such fluctuating biomolecular condensates at the required resolution in space and time, an atomic-level picture that connects the structure and dynamics of protein condensates with the molecular thermodynamics is largely lacking. Atomistic molecular dynamics (MD) simulations with explicit solvent can, in principle, provide such detailed insights into the structure, composition, and dynamics of protein condensates. However, the huge computational cost renders it impossible to access the slow timescales needed to observe the process of protein condensate formation. In addition, very large simulation systems are required to accommodate the two coexisting phases, which further increases the computational effort. To the best of our knowledge, there have been only few atomistic MD simulation studies of biomolecular condensates, ^56–60^ none of which simulated the actual process of phase separation. Phase separation processes, however, have been simulated using coarse grained descriptions that are computationally efficient and greatly accelerate the simulations.^49,61–65^ However, such coarse grained models have limitations concerning their ability to provide accurate, atomically-detailed thermodynamic and dynamic evaluations of the protein and, in particular, of the solvent, which is the scope of the present work.

Experimentally, protein condensates are challenging to study with high resolution. One particular challenge also includes the accurate measurement of the protein concentration in the condensate. McCall et al. ^66^ have used quantitative phase microscopy to determine the condensate concentration of a full-length FUS protein (tagged with a fluorescent protein) to be 337 ± 8 mg mL^−1^. Murakami et al. have recently used Raman imaging microscopy to determine the concentration of proteins in droplets formed as a result of LLPS.^67^ This method has also been applied on FUS-LCD, yielding condensate and dilute phase concentrations of a recombinant FUS-LCD of 15 mM (320 mg mL^−1^) and ∼60 *µ*M (1 mg mL^−1^), respectively, under the experimental conditions (room temperature, physiological salt concentration).^48^ Another study by Murthy et al. ^55^ reported the condensate concentration to be 477 mg mL^−1^ for FUS-LCD. The concentration range obtained from the above experiments is quite broad, but they nevertheless provide a useful concentration window that would represent a FUS condensate.

As simulating the actual process of phase separation is currently impossible with atomistic MD simulations, knowing the concentrations of the dilute and dense phases of FUS-LCD is required for extracting the thermodynamic contributions underlying condensate formation from MD simulations. Since free energy is a state function, separate simulations of (a set of) homogeneous FUS-LCD solutions, including the final protein condensate concentration (Figure 2), and of the dilute phase yield access to the states necessary for computing the thermodynamic changes associated with the process. For the FUS-LCD condensate concentration, we used a concentration of 350 mg mL^−1^. For the dilute phase we consider bulk water because the concentration of FUS-LCD is very low (1 mg mL^−1^, see above).^48^

**Figure 2:**
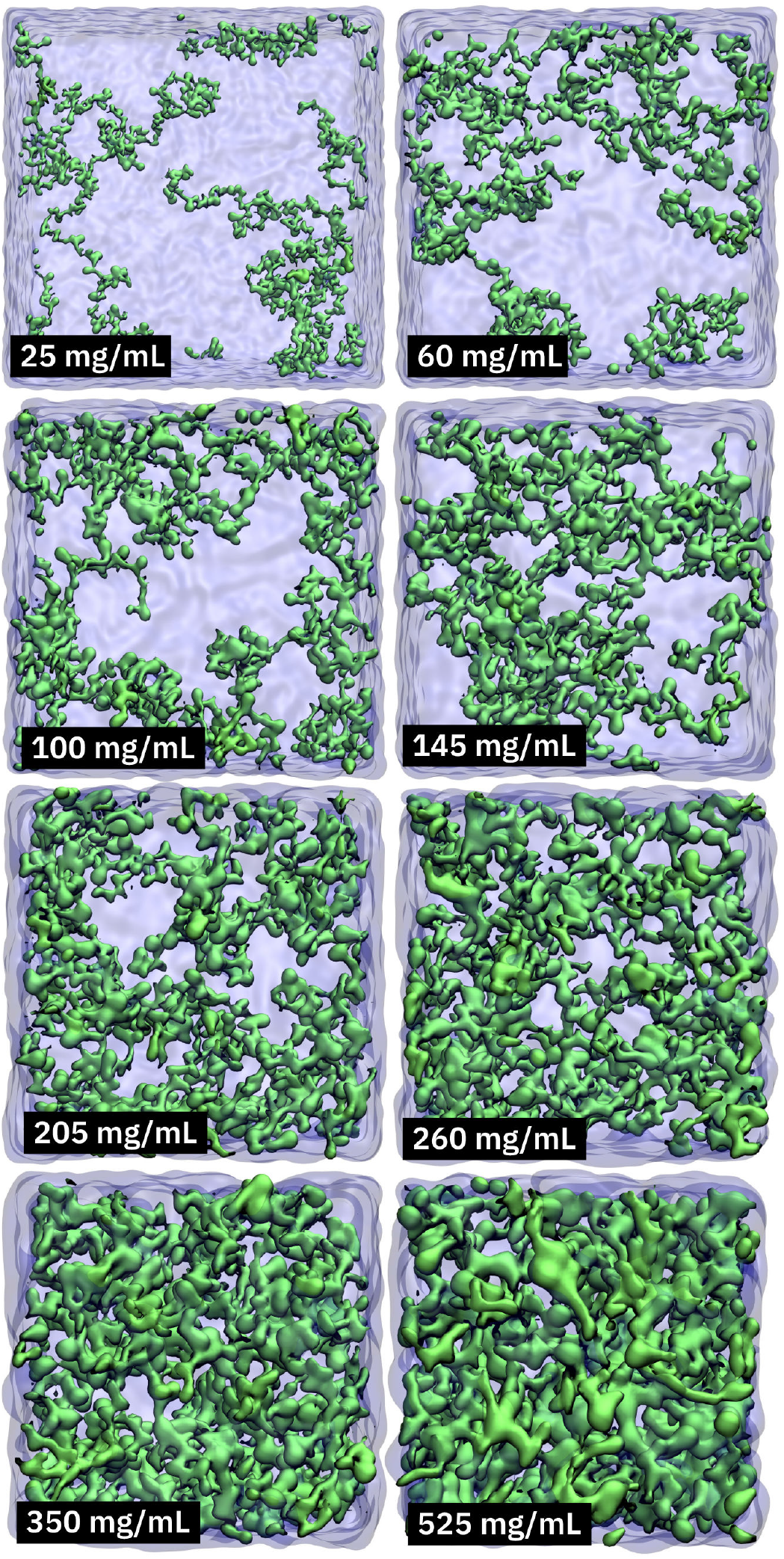
Snapshots of simulated systems. Each of the simulated systems contains eight FUS-LCD molecules (residues 1 - 163 of human FUS, UniProt ID P35637).

Here, we unravel the thermodynamic driving forces underlying the formation of FUSLCD condensates using atomistic MD simulations, which enable the decomposition of the total free energy change into contributions from protein–protein, protein–water, and water– water interactions. The analyses provide insights into the entropy and enthalpy changes associated with each of these components. A tug-of-war between the entropy of the released and the retained water molecules is found to be a major mechanistic determinant in the process. In addition, favorable protein–protein interactions also contribute substantially to the thermodynamic driving forces. Entropy-dominated solvation effects and enthalpy-dominated protein interactions are found to be comparable in magnitude, supporting the notion that for a complete and quantitative picture, both effects need to be considered.

## Results and Discussion

### Solvent rearrangements upon condensate formation

Protein condensates are formed via LLPS when the concentration reaches a threshold value (which can vary depending on conditions such as temperature, pressure, and other external factors). Upon condensation, the protein molecules are brought into close proximity to each other until eventually, at high concentrations, the protein hydration layers (PHLs) start to overlap and are eventually even partially stripped off. Consequently, the properties of the water that is confined inside a dense condensate are expected to be affected and to be distinctly different compared to the dilute solution. As shown in Figure 1, some of the waters are released from the condensate into the dilute phase, while others remain inside the droplets.

To characterize the changing hydration water populations upon condensate formation, the number of water molecules within 0.3 nm from the protein surface 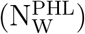 was analyzed, that is, we focused on the water molecules in the first hydration shell that are in direct contact with the protein. In Figure 3a the number of hydration waters per protein molecule is plotted as a function of the protein concentration. The numbers in this plot are normalized with respect to the number of water molecules in the hydration layer of a simulation system with only a single FUS-LCD molecule, that is, under “infinite dilution” conditions. Even at 25 mg mL^−1^, the PHL has 6% fewer water molecules than at infinite dilution. As expected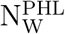, decreases with increasing protein concentration. The decrease in hydration water population is due to the release of some water molecules from the PHL into the dilute phase in the course of condensation. However, the proteins in the condensate retain a substantial fraction of their first shell PHL water molecules even at high concentrations.

**Figure 3:**
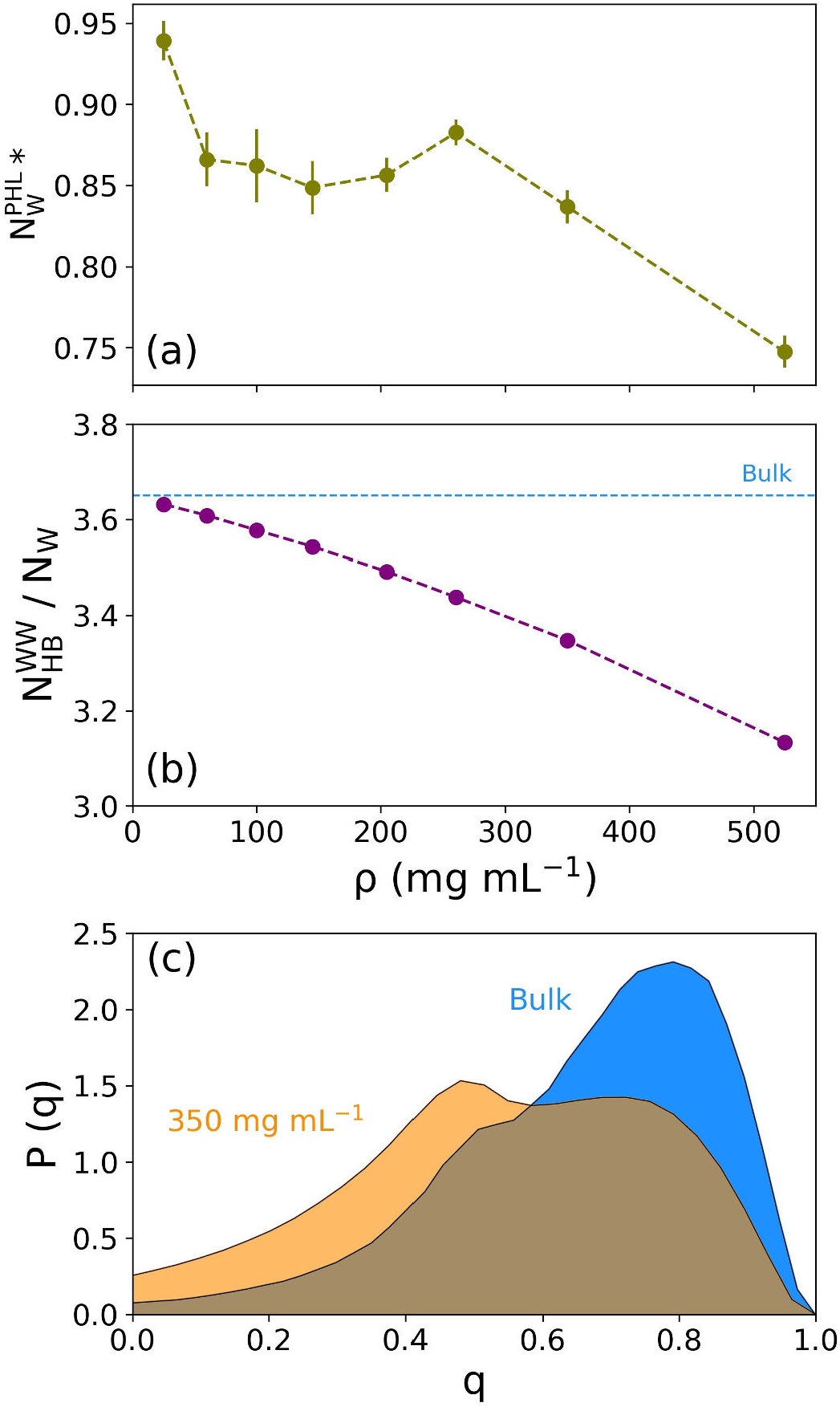
Hydration properties at different FUS-LCD concentrations. (a) Number of water molecules in the protein hydration layer (defined here as water molecules within 0.3 nm of the protein surface). 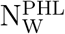 is plotted against the protein concentration *ρ*. The asterisk (*) denotes that the numbers are normalized with respect to the number of hydration waters found for a single FUS-LCD protein in the high-dilution limit. (b) The number of water– water hydrogen bonds per water molecule is plotted as a function of *ρ*. (c) Tetrahedral order parameter distribution of water in 350 mg mL^−1^ FUS-LCD solution (orange) compared to bulk water (blue). Data are presented as mean values ± standard deviation (SD) over three repeat simulations (the statistical errors in (b) are smaller than the size of the dots).

The spatial confinement imposed by the high protein concentration affects the hydrogen bond (HB) network between the water molecules. With increasing protein concentration, perturbations in the HB network result in the loss of water–water HBs. Besides the fact that the water concentration itself is decreased in the condensates, the waters that are retained inside the condensate form HBs with the protein. This results in a decrease in the number of water–water HBs formed per water molecule (Figure 3b). This spatial confinement effect is also reflected in the tetrahedral arrangement of water molecules, with decreasing population of water molecules in a tetrahedral local environment with increasing *ρ*. The local environment around a water molecule can be quantified by the tetrahedral order parameter (Eq. 1)^68–70^

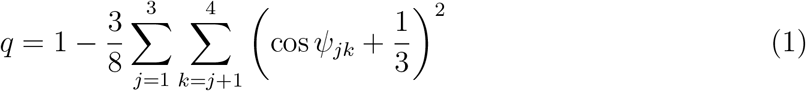

where *ψ*_*jk*_ denotes the angle between the *j*^*th*^, the central, and the *k*^*th*^ water molecule. For a perfect tetrahedral arrangement, *q* = 1, whereas *q* = 0 for a completely non-tetrahedral configuration. A comparison between the distributions of *q* for bulk water and for water in the FUS-LCD condensate (350 mg mL^−1^) shows that the peak around *q* = 0.8 decreases in the concentrated protein solution as compared to bulk water, concomitant with an increase in the peak intensity around *q* = 0.5 (Figure 3c). Thus, the water molecules retained in the protein condensates experience a less tetrahedral local environment and form more twodimensional trigonal arrangements to accommodate intermolecular hydrogen bonding.

The change in the population of water surrounding the proteins, as analyzed above, classifies the water molecules in the system into two distinct categories, i.e., *released* and *retained*. While the former are transferred into the dilute (bulk-like) phase, the latter face a more confined surrounding inside the condensate. To understand the putative thermodynamic consequences of these changes in water populations, we next focus on the tug-of-war between the released and the retained water molecules in terms of their enthalpic and entropic contributions to the free energy. To that end, we first quantify the number of the released and retained water molecules.

Starting from a (hypothetical) homogeneous solution with a protein concentration *ρ*, the number of released water molecules is given by the difference in the number of waters in the initial system and in the condensate (Eq. 2)

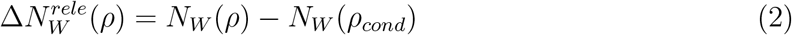

Eq. 2 gives the number of water molecules that would be released into the dilute phase if the protein concentration in the initial homogeneous system (that is, prior to phase separation) was *ρ* (see Figure 1). Since *ρ* = *ρ*_*cond*_ in the condensate, 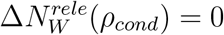

In contrast to the released waters, the number of water molecules that are retained in the condensate is constant (Eq. 3),

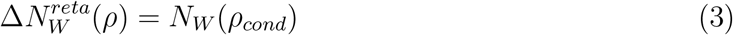

assuming that the condensate has a constant (fixed) concentration. This is shown graphically in Figure 4 and the corresponding numbers are given in Table S1.

**Figure 4:**
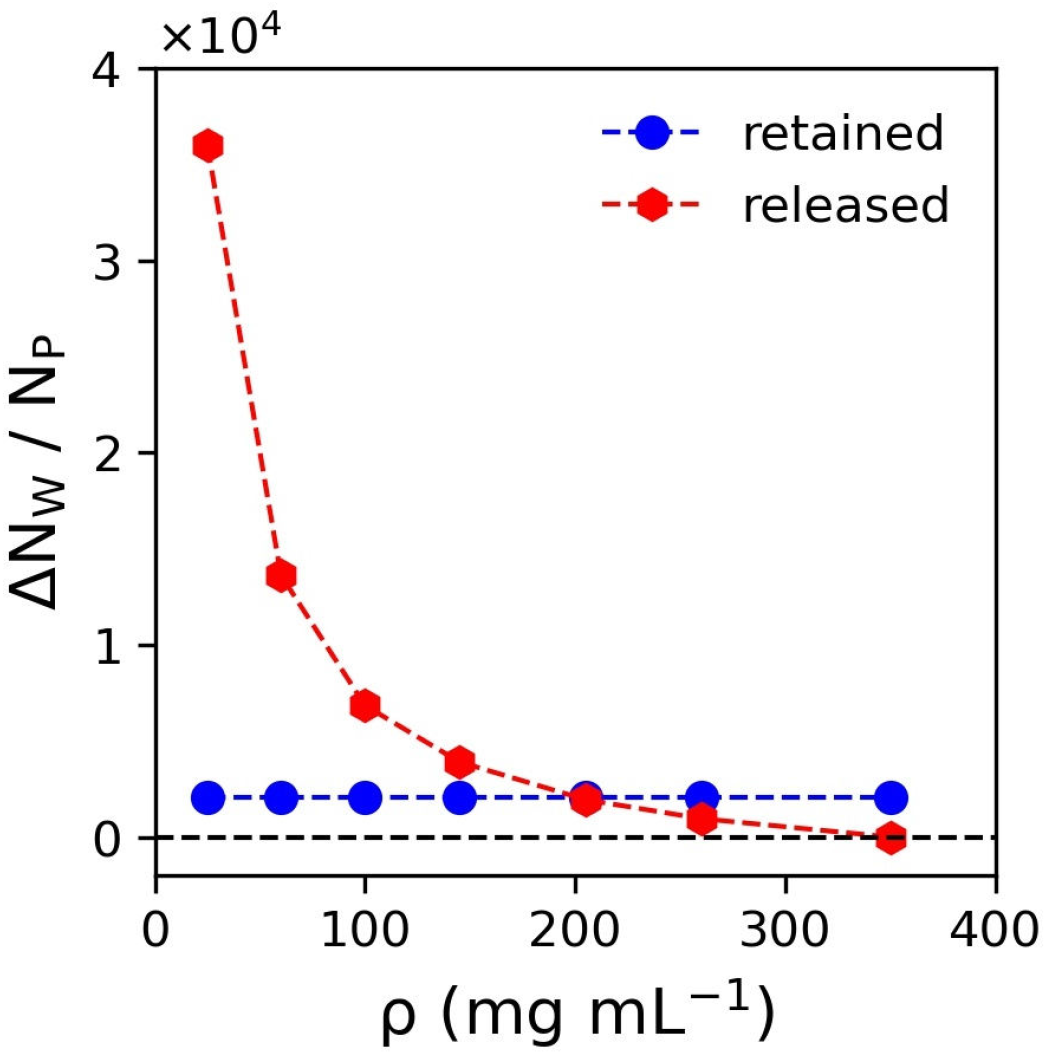
Number of released and retained water molecules per protein. The number of released molecules decreases (red dashed line) as the FUS-LCD concentration *ρ* approaches the condensate concentration of 350 mg mL^−1^, where every protein is solvated (on average) by 2064 water molecules (blue dashed line), which are referred to as the retained waters.

### Thermodynamic signatures of FUS-LCD condensate formation

Knowing the changes in solvent populations during the FUS-LCD condensation forms the basis for understanding the underlying thermodynamic driving forces. The total free energy change (∆*G*) upon condensation can be defined to have two contributions (Eq. 4)

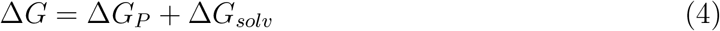

One part of the free energy change is directly governed by the proteins themselves (∆*G*_*P*_), which includes changes in the protein enthalpies (∆*H*_*P*_) and entropies (∆*S*_*P*_)

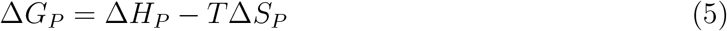

∆*H*_*P*_ comprises of the intra- (∆*E*_*intra*_) and intermolecular (∆*E*_*inter*_) potential energies of the proteins present in the system (Eq. 6), and hence, refers to the total protein–protein interactions (∆*E*_*PP*_). Since the systems are hardly compressible, the *p*∆*V* term can be neglected and the enthalpy difference be accurately approximated by the interaction energy difference,

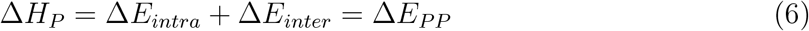

These energy terms are readily available (within statistical error limits) from the force field interaction energies, averaged over the MD trajectories.

The protein entropy in Eq. 5 is very challenging, particularly for large and flexible molecules like IDPs. The entropic penalty paid by the proteins upon the formation of condensates has two parts. First, the proteins experience an entropy penalty that is associated with the restriction of the overall translational and rotational motions of the proteins in the condensates. These two contributions are hard to separate from each other for IDPs, because their high degree of flexibility renders it challenging to decouple overall motions from internal dynamics. However, the free energy change due to the entropy of (de)mixing can be simply estimated from the concentration ratio, ∆*G* = *RT* ln(*ρ*_*cond*_*/ρ*_*dil*_). With the condensate and dilute phase values of 350 mg mL^−1^ and 1 mg mL^−1^, respectively, one obtains ∼14.5 kJ mol^−1^ for the associated free energy penalty. This contribution was neglected because it is much smaller than the other thermodynamic contributions to the condensate formation (and also much smaller than the statistical uncertainties due to limited sampling in the MD simulations), see below.

By construction, the setup of our simulation systems as homogeneous FUS-LCD solutions (with different concentrations) neglects the energy associated with the formation of an interface upon phase separation. Biomolecular condensates typically have ultralow surface tensions^71^ ranging from *γ* = 0.0001 mN/m up to ca. 0.5 mN/m (for comparison, the air/water surface tension is 72 mN/m at 298 K). To roughly estimate the energetic penalty of interface formation, we calculated the interface free energy of spherical FUS-LCD droplets (c = 350 mg mL^−1^) with radii from 20 nm up to 5000 nm, thus covering a broad size range. The resulting free energy penalties per protein molecule are small, with the maximum value of 4 kJ mol^−1^ for a (hypothetical) very small nanodroplet with 20 nm radius and *γ* = 0.5 mN/m (which is a factor of more than 150 larger than the actual surface tension of 0.003 mN/m reported for FUS^72^). Therefore, this contribution is neglected in the present work.

Furthermore, one needs to consider the conformational entropy change (∆*S*_*conf*_) that arises from the differential restriction of the conformational flexibility (and the corresponding changes in the configuration space densities) of the proteins in the condense phase compared to the dilute solution. As is discussed in more detail below, the protein conformational entropy change is notoriously hard to quantify due to sampling limitations, but our estimate indicates that the −*T* ∆*S*_*conf*_ contribution to the free energy change upon FUS-LCD condensate formation is much smaller than the other, more significant contributions related to solvation and protein–protein interactions.

The second part of the total free energy (Eq. 4) comes from solvation

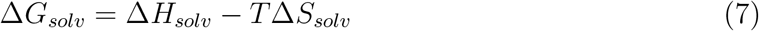

Note that the index “solv” usually indicates the solvation process defined as the transfer of a solute from the vacuum to the solution, and hence in the present case, where the difference in solvation between protein solutions of different concentrations is considered, the notation ∆∆*G*_*solv*_ (or accordingly also for the enthalpy and entropy) might be considered more appropriate. However, for simplicity and consistency, we here denote the changes with a single ∆.

Both the solvation enthalpy and entropy in Eq. 7 can be formally divided into protein– water and water–water contributions

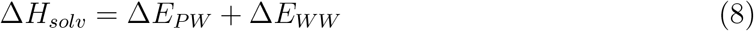

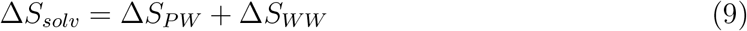

The enthalpy and entropy terms involving water–water interactions exactly cancel each other (Eq. 10)

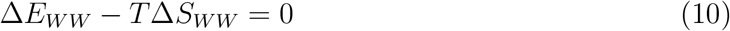

as originally proposed by Ben-Naim ^73^ and later shown by Yu and Karplus ^74^ and by Ben-Amotz.^75^ Recently, Heinz and Grubmüller clarified the importance of the precise definition of the solvent–solvent interactions in this context.^76^ From Eqs. 7-10 it follows that the total solvation free energy depends only on the protein–water terms

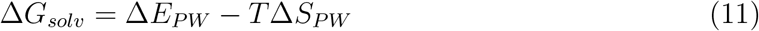

Like in the case of protein–protein interaction energies, ∆*E*_*PW*_ can be directly calculated from the force field energy terms, averaged over the simulation trajectories. However, the entropy of water is more difficult to evaluate due to the diffusive nature of the liquid. We used the 2PT method introduced by Lin et al. ^77^ for that purpose (see Methods).

Figure 5a shows that the total molar entropy (*S*_*tot*_) of water is decreasing with increasing protein concentration. At higher protein concentrations, the water molecules are located in an increasingly crowded and confined environment that strongly attenuates their configuration space, resulting in lower entropy. A cross-over is observed at around *ρ* = 150 mg mL^−1^ from one linear gradient to another, steeper one (see also Figure S1). The underlying picture becomes clearer when the total entropy is decomposed into its translational (*S*_*tr*_) and rotational (*S*_*rot*_) contributions (Figure 5b). Here, the entropies are normalized with respect to their bulk values (Table 1).

**Table 1:** Entropy of bulk water (a99sb-disp water model) evaluated using the 2PT method. The errors denote standard deviation.

| Entropy<br>term | Value<br>(J mol<sup>-1</sup> K<sup>-1</sup>) |
| --- | --- |
| $S_{tot}$ | $54.7 \pm 0.2$ |
| $S_{tr}$ | $43.8 \pm 0.2$ |
| $S_{rot}$ | $10.9 \pm 0.1$ |

**Figure 5:**
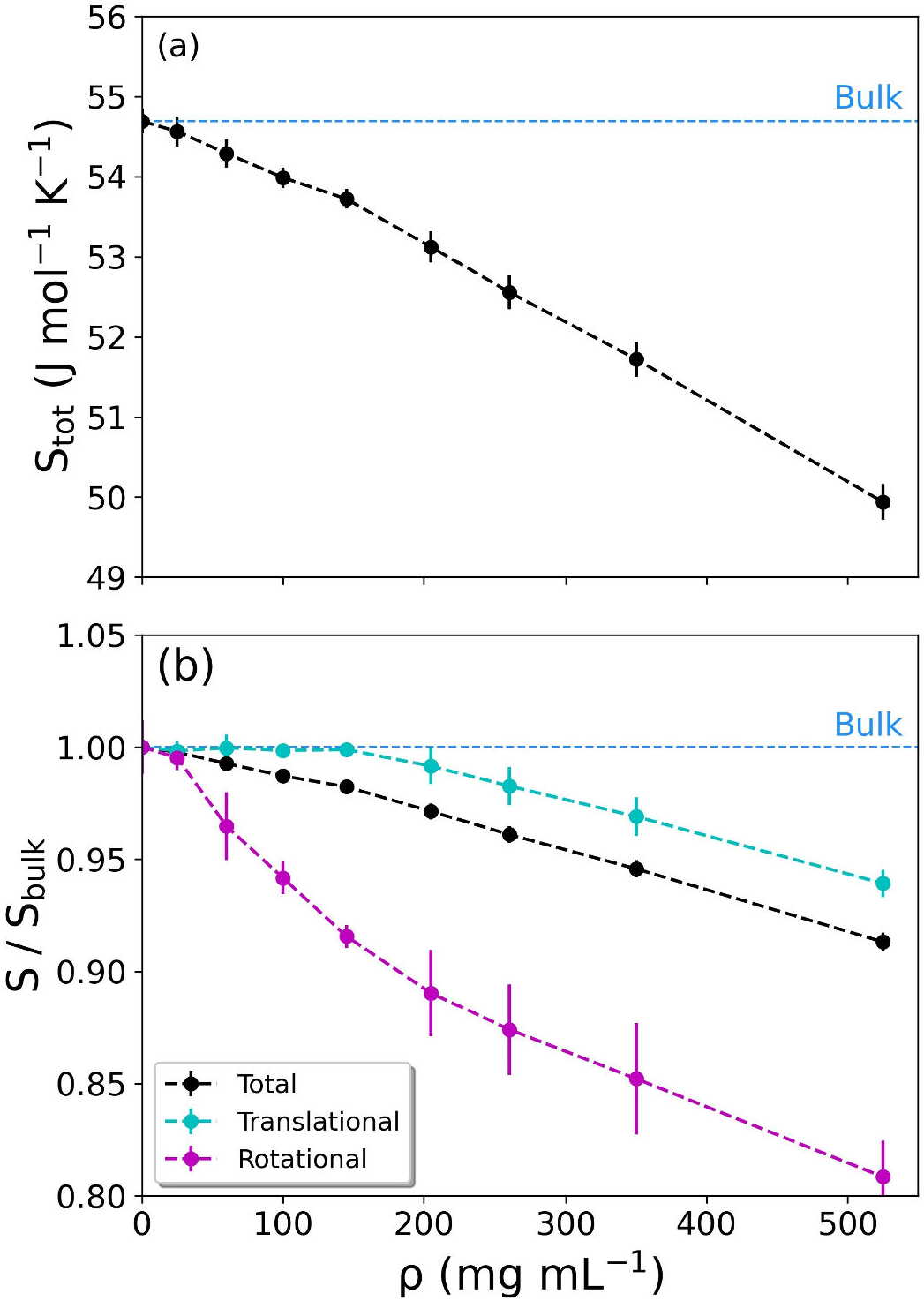
Entropy of water as a function of protein concentration (*ρ*). (a) Total entropy. The horizontal dashed line represents the bulk water value. (b) Decomposition of the total entropy (black curve) into translational (cyan) and rotational (magenta) contributions. The values in (b) were normalized with respect to bulk water. Data in panels (a) and (b) are presented as mean values ± standard deviation (SD) over three repeat simulations.

Figure 5b shows that the cross-over between the two linear regimes has its roots in the differential changes of the translational and rotational entropy components with *ρ. S*_*tr*_ remains almost unchanged up to *ρ* ∼150 mg mL^−1^, after which it starts to drop linearly. In contrast, *S*_*rot*_ starts to decrease already at low protein concentrations and faces a cross-over at *ρ* ∼150 mg mL^−1^, thereafter continuing to decrease with a slope that is ∼2.4 times smaller than at low *ρ*. Hence, it seems that the entropy linked to the rotational motions of water is much more susceptible to the presence of proteins. Moreover, the crossover concentrations of *S*_*tr*_ and *S*_*rot*_ appear to coincide. A deeper investigation of the underlying effects is intriguing, but is out of the scope of the present work.

To scrutinize the accuracy of the 2PT method, the free energy perturbation (FEP) method^78,79^ was used to compute the entropy of bulk water from the free energy of solvation and enthalpy of vaporization. The total entropy from FEP is 54.4 J mol^−1^ K^−1^, which is very close to the value of 54.7 J mol^−1^ K^−1^ obtained from 2PT (Table 1). A detailed description of the calculations is given in the SI. FEP does not yield the translational and rotational entropies separately, but it is based on a rigorous statistical mechanics framework (that does not rely on an assumed decomposition of the density of states into a solid-like and a gas-like part), and therefore FEP can provide a meaningful and independent validation of the entropy estimates obtained with 2PT.

The property of interest here is the entropy change (∆*S*) associated with the formation of a FUS-LCD condensate. The release of water into the dilute surrounding is associated with an increase in entropy that can be calculated by subtracting the entropy of water in a system with protein concentration *ρ, S*(*ρ*), from that of bulk water, *S*(*ρ*_*dil*_), which we take as a reference for the dilute phase because the concentration of FUS-LCD in the dilute phase is very low, about 1 mg mL^−1 48^). The total entropy change is then obtained by multiplying with the number of released waters (Eq. 2). Similarly, the total entropy penalty associated to water retention can be computed by subtracting the entropy of water in a system from that in the protein condensate phase (*ρ*_*cond*_ = 350 mg mL^−1^), multiplied by the number of retained water molecules. This is shown in Eq. 12.

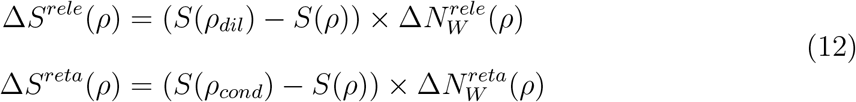

Note that the numbers of released and retained waters in Eq. 12 are normalized with respect to the number of protein molecules (see Figure 4). To obtain the total solvent entropy change, the two contributions are added (Eq. 13)

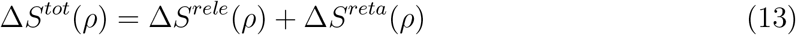

∆*S*^*tot*^(*ρ*), however, is not effective in the overall thermodynamics of the process because the part of it originating from water–water interactions is canceled out by a compensating enthalpy term (Eq. 10). Hence, the entropy bill obtained from Eq. 12 only comprises of the non-canceling term, *T* ∆*S*_*PW*_, which can be obtained by subtracting the calculated water– water interaction energy change from the total solvation entropy term (Eq. 14).

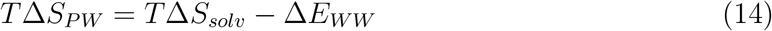

The other part of free energy originates from the interaction energies. Figure 6 plots the *ρ*-dependence of protein–protein (*E*_*PP*_), protein–water (*E*_*PW*_), and water–water (*E*_*WW*_) interaction energies, normalized by the corresponding numbers of molecules.

**Figure 6:**
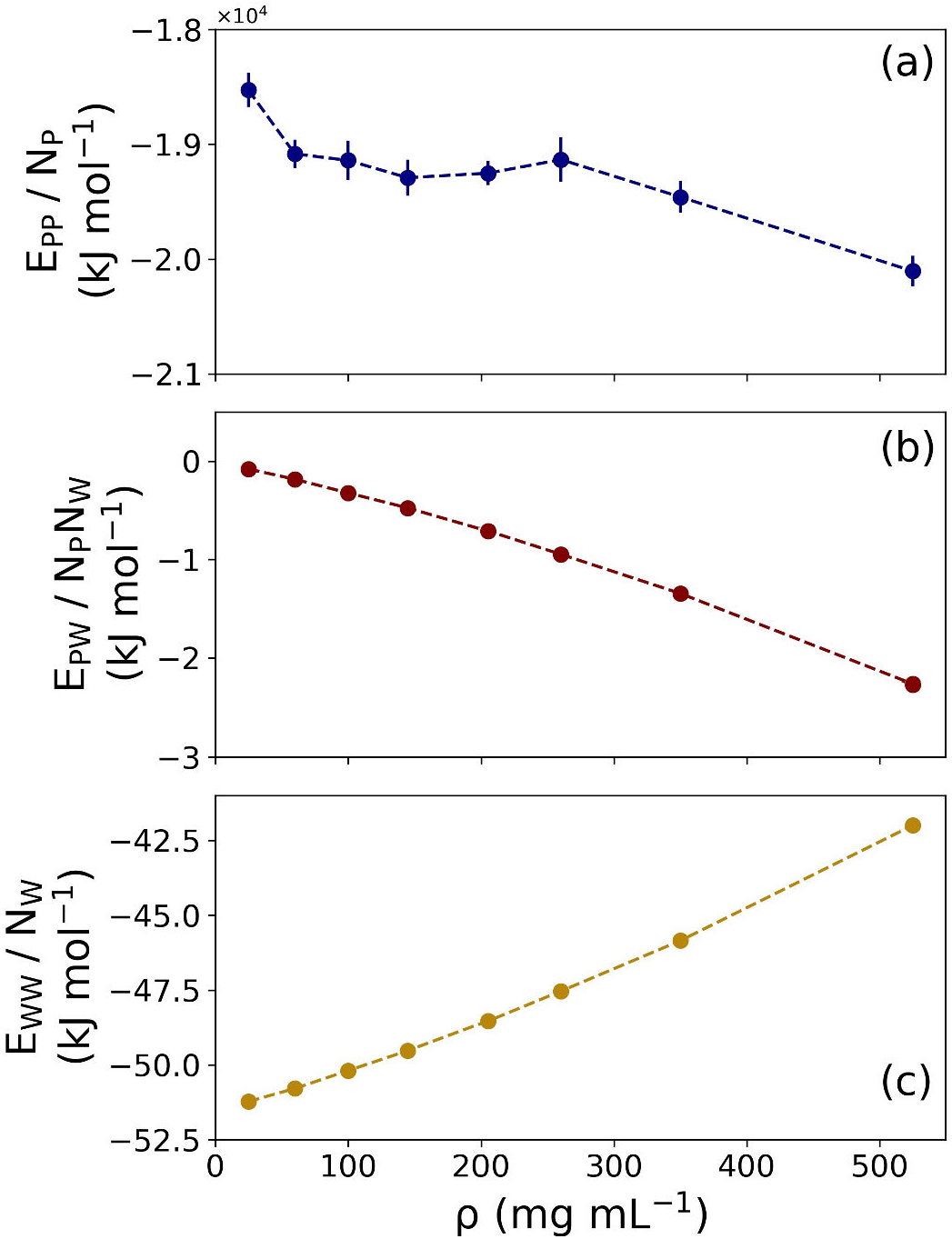
Interaction energies between the different components as a function of protein concentration. (a) Protein–protein interaction energy (per protein molecule), including intra- and inter-protein contributions. (b) Protein–water interaction energy (per water molecule). (c) Water–water interaction energy (per water molecule). Data in panels (a), (b), and (c) are presented as mean values ± standard deviation (SD) over three repeat simulations (the statistical errors in (b) and (c) are smaller than the size of the dots).

Figure 6a shows that *E*_*PP*_ decreases with increasing *ρ*. This is a result of favorable protein–protein interactions in the condensate, as have been reported before in experimental and theoretical studies. ^17,27,80^ The favorable change in *E*_*PW*_ (Figure 6b) arises from the increased fraction of protein–water interactions, mostly through hydrogen bonds, with increasing protein concentration. The simultaneous decrease of water–water HBs (Figure 3b) accounts for the destabilization of water–water interactions, which results in the increase of *E*_*WW*_ with *ρ* (Figure 6c).

In addition to entropy, another major thermodynamic contribution is associated to the changes in the interaction energies upon FUS-LCD condensate formation. The above considerations for calculating entropy changes of released and retained water molecules (Eq. 12) also apply to the changes in energies (Eq. 15)

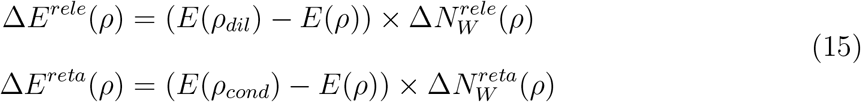

Each of the terms in Eq. 15, ∆*E*^*rele*^(*ρ*) and ∆*E*^*reta*^(*ρ*), has two contributions, which are the canceling water–water (∆*E*_*WW*_) and the non-canceling protein–water (∆*E*_*PW*_) interaction energies. The total change in solvation enthalpy (∆*H*_*solv*_) is the sum of these two terms (Eq. 8), although the former term does not influence the change in free energy due to cancellation (Eq. 10).

The scenario is somewhat different for the changes in protein–protein interaction energies, ∆*E*_*PP*_ . Since our reference dilute phase is bulk water (see above), in our analyses we assume that proteins are present only in the condensate phase. Consequently, there are no “released proteins” upon condensate formation and therefore, the total change in protein–protein interaction energy is entirely determined by the protein interactions in the condensate (Eq. 16)

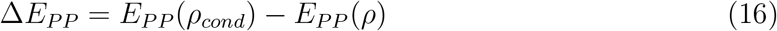

The changes in the thermodynamic quantities discussed thus far are plotted in Figure 7, with separate contributions from retained water, released water, and the total change resulting from the sum of these two individual components (the “tug–of–war”). Figure 7a shows the total solvation entropy change (Eq. 9, see also Table S2). The entropy change for released water favors condensate formation because water molecules explore a larger configuration space in the bulk-like dilute phase. At the same time, the retained water molecules are in a more crowded environment, and thus have a decreased entropy that disfavors phase separation. In this entropy tug-of-war, the contribution of the released water molecules slightly outweighs that of the retained waters, resulting in a slightly negative −*T* ∆*S*_*solv*_ (Figure 7a, inset), and hence the solvation entropy change favors condensate formation.

**Figure 7:**
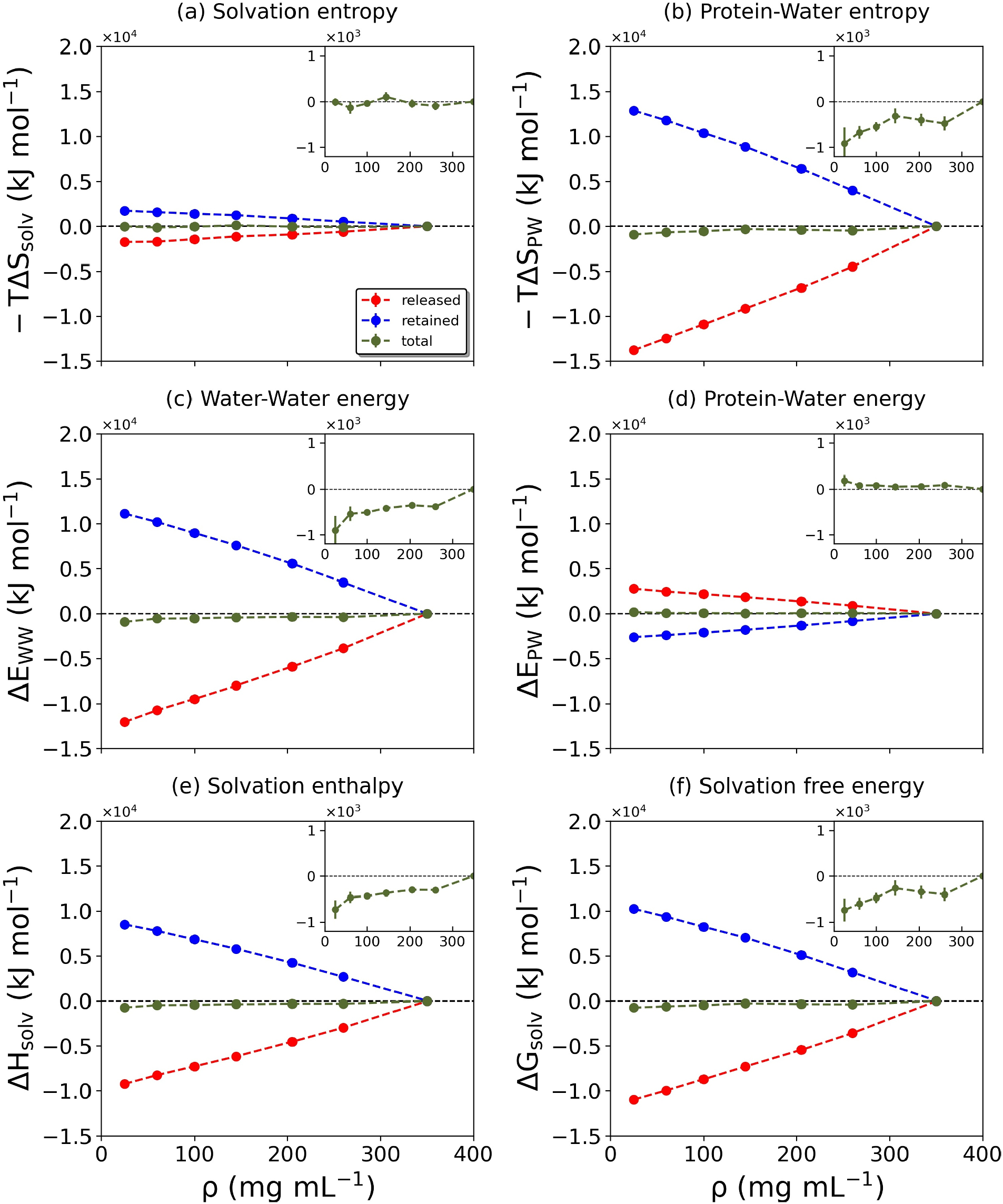
Changes of the solvation-related thermodynamic quantities. The quantities plotted in (a)-(f) are indicated at the top of each panel. The dashed red, blue, green lines denote the released, retained, and total water contributions, respectively. Data are presented as mean values ± standard deviation (SD) over three repeat simulations.

The entropic driving forces become clearer from the noncanceling entropy changes originating from the protein–water interactions, Eq. 14 (Figure 7b and Table S3). Although the thermodynamic contributions from released and retained water changes compensate each other to a large extent, the total change in −*T* ∆*S*_*PW*_ remains favorable (Figure 7b, inset) and substantially greater than −*T* ∆*S*_*solv*_. The difference in the magnitudes of −*T* ∆*S*_*solv*_ and −*T* ∆*S*_*PW*_ originates from the canceling water–water term, ∆*E*_*WW*_ = *T* ∆*S*_*WW*_, shown in Figure 7c and Table S4. This ∆*E*_*WW*_ contribution is larger than the noncanceling ∆*E*_*PW*_ energy term (Figure 7d and Table S5).

Interestingly, the released and retained waters behave oppositely in the water–water and protein–water interaction energy contributions (Figure 7c and d), with the former (water– water) term dominating the total solvation enthalpy change (Figure 7e and Table S6). On approaching the condensate concentration, an increasing fraction of the retained water molecules form protein–water HBs at the cost of water–water HBs (Figure 3b). Hence, the interaction energy of the retained waters with the proteins becomes more favorable. Simultaneously, the decrease in the water–water HBs leads to the increase of the interaction energy, that is, unfavorable ∆*E*_*WW*_ . At the same time, the released waters gain more water–water contacts at the expense of losing contacts with proteins.

Figure 7f shows that the change in total solvation free energy is favorable for the formation of FUS-LCD condensates (see also Table S7). Hence, in the described “tug-of-war” between retained and released water molecules, the latter win against the former, resulting in a significant net thermodynamic driving force for the formation of FUS-LCD condensates.

The other part of the total free energy change is not linked to solvation but originates from the proteins. As discussed above in the context of in Eq. 16, this energy contribution is entirely governed by the protein interactions in the condensate (Figure 8 and Table S8).

**Figure 8:**
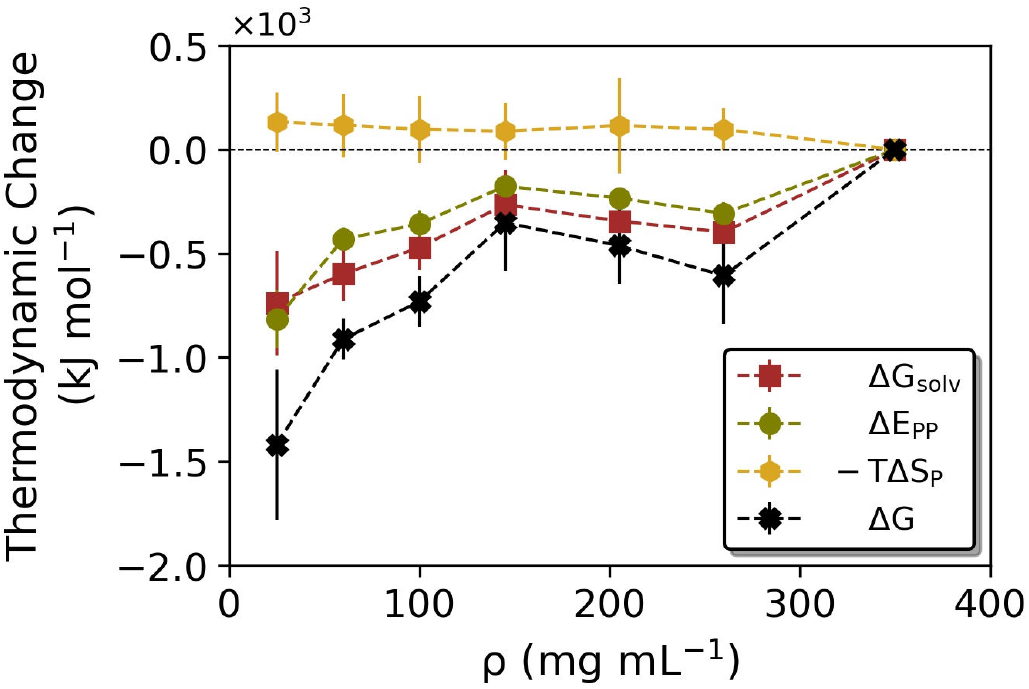
Thermodynamic driving forces of FUS-LCD condensate formation. The contributions from total solvation free energy (∆*G*_*solv*_, brown dashed line), protein–protein interaction energy (∆*E*_*PP*_, green dashed line), and protein conformational entropy (∆*S*_*P*_, orange dashed line) are plotted together with the resulting total free energy change upon formation of a FUS-LCD condensate (∆*G* = ∆*G*_*solv*_ + ∆*E*_*PP*_ − *T* ∆*S*_*P*_, black dashed line), starting from a (hypothetical) homogeneous solution with protein concentration *ρ*. Data are presented as mean values ± standard deviation (SD) over three repeat simulations.

The protein–protein interaction energy, ∆*E*_*PP*_ is attractive at higher concentrations due to favorable contacts between the FUS-LCD molecules. This finding is in line with the fact that FUS-LCD has a large fraction of polar residues, but does not carry a large net charge (-2 in our simulations) that would result in repulsive Coulomb interactions at high concentrations.

Finally, the missing thermodynamic contribution is the change in protein conformational entropy upon condensate formation (Eq. 17)

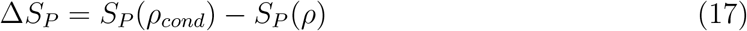

which is expected to be unfavorable due to the increased confinement of the proteins. As explained in Methods, the protein conformational entropy was estimated using a nearest-neighbor method.^81^ Figure 8 (orange curve, see also Table S9) shows that indeed, −*T* ∆*S*_*P*_ is unfavorable, but it is much smaller than the solvation free energy and protein–protein interaction energy. As a check, additional control simulations were performed in which two trajectories, one for the 25 mg mL^−1^ system and one for the 350 mg mL^−1^ system, were extended to 400 ns and the protein conformational entropies were recalculated based on the final 300 ns of these trajectories. The resulting −*T* ∆*S*_*P*_ value of 87.3 kJ mol^−1^ lies within the error bounds plotted in Figure 8. The relatively small difference in protein conformational entropy between the dilute and dense phases is in line with the distributions of the radii of gyration of the individual FUS-LCD polypeptide chains in the simulation systems at 25 and 350 mg mL^−1^ (Figure S2), which suggest that a pronounced degree of flexibility is retained also within the condensate.

The accurate estimation of (changes in) protein conformational entropy is notoriously hard because it requires exhaustive conformational sampling, ^82,83^ which is impossible with all-atom MD for an IDP of this size. In light of these limitations, we consider ∆*S*_*P*_ to be a rough estimate. However, the main finding that ∆*G* is favorable is robust with respect to uncertainties in the protein conformational entropy estimate, as changing −*T* ∆*S*_*P*_ even by one order of magnitude does not change the sign of the total free energy change.

Combining all these contributions, the total free energy change is negative (black curve in Figure 8, see also Table S10), in agreement with the experimentally observed spontaneous LLPS of FUS-LCD. The two largest thermodynamic driving forces that favor condensate formation are the noncanceling (protein–water) solvation entropy and the protein–protein interaction energy. These two contributions are comparable in magnitude, showing that solvation effects are as important as protein interactions for the formation of FUS-LCD condensates.

Finally, since there is no general consensus on the exact FUS-LCD concentration in the condensate, we repeated our analyses using a condensate concentration of 525 mg mL^−1^ instead of 350 mg mL^−1^ as a reference. The results are shown in Figures S3-S5 and confirm the above conclusions, in that the thermodynamic profiles look very similar. The magnitudes of the individual contributions are larger, as expected due to the more pronounced concentration difference between the condensate and the dilute phase.

To summarize this work, we used large-scale atomistic MD simulations to analyze in molecular detail the thermodynamic driving forces underlying the condensate formation of the intrinsically disordered low complexity domain (LCD) of the human fused in sarcoma (FUS) RNA-binding protein. An understanding of the individual contributions of proteins and water is provided, ultimately yielding a complete thermodynamic picture.

The formation of protein condensates leads to a fraction of the hydration water molecules being released into the dilute phase, which in the case of FUS-LCD has a very low protein concentration and is bulk water-like. The remaining solvent fraction is retained inside the protein condensate, where it is strongly confined by the dense protein environment. The tug-of-war between these two categories of water molecules determines the solvent thermodynamics of the overall process. The retained waters have substantially lower entropy. Furthermore, the hydrogen bonds formed between the retained water molecules and the proteins perturb the extended hydrogen bond network among the water molecules themselves and weaken the water–water interactions. In contrast, the released waters in the dilute phase loose interactions with the proteins and form more water–water H-bonds, and they gain entropy. This latter contribution dominates, and thus the released waters win the tug-of-war, resulting in an entropy-dominated favorable solvation force for FUS-LCD condensate formation.

Favorable protein–protein interaction energy also significantly contributes to the FUS-LCD condensate formation, comparable in magnitude to the total solvation free energy change. This shows that for FUS-LCD, both water and protein are equally significant in the thermodynamics of protein condensate formation via LLPS.

By construction, the present study focuses on the formation of FUS-LCD condensates *in vitro*, that is, aqueous two-phase systems in which the dense condensate is surrounded by a dilute phase. Of course, the actual scenario inside a biological cell is much more complicated. The cell cytoplasm is crowded, and the surrounding of a condensate in a cell, and hence the environment that the released water molecules enter, is far from dilute. Furthermore, the condensate will typically not be comprised only of a single (protein) species, but of multiple different components, including RNA, which can have internal structural organization.^84,85^ Therefore, the magnitudes of the thermodynamic contributions identified in this work should be interpreted rather as upper limits, especially the changes associated with the released water. Nevertheless, we expect that the insights and overall conclusions of this work are transferable to biological cells, at least at a qualitative level, as long as there are distinct regions with different concentrations.

## Methods

### Setup of simulation systems

All simulations were performed using the GROMACS (version 2020.1) molecular dynamics simulation package.^86,87^ The the Low Complexity Domain (LCD) of the human Fused in Sarcoma (FUS) RNA-binding protein (UniProt ID: P35637) was simulated, which encompasses residues 1 to 163 (out of a total of 526 residues) and has a molar mass of 17.2 kDa. FUS-LCD is an intrinsically disordered protein (IDP) domain for which no experimental high-resolution structure is available.

The initial atomic coordinates of unfolded FUS-LCD were generated with AlphaFold,^88,89^ which predicts an extended conformation without any secondary structure elements. The first aim was to obtain pre-equilibrated systems at the desired protein concentrations. To accelerate this step, coarse-grained (CG) MD simulations were carried out using the Martini 2.2^90,91^ force field. The CG-Martini topology, as generated with the martinize.py script available on the martini website (www.cgmartini.nl), did not contain any elastic network bonds between CG beads. We inserted 8 copies of the coarse-grained FUS-LCD in a cubic periodic simulation box with ∼27 nm edge length, corresponding to a protein concentration of ∼10 mg mL^−1^. 176366 CG water beads and 16 sodium ions (for neutralization of the charge of the simulation box) were added to the system (1 CG water bead effectively represents 4 water molecules in the Martini force field). Protein–protein Lennard-Jones (LJ) interactions were scaled according to the protocol of Stark et al.,^92^ which was recently used by Benayad et al. for studying FUS-LCD condensate formation.^49^ The well depths of the LJ 6,12 potential were modified with a scaling parameter *α* according to *ϵ*_*α*_ = *ϵ*_0_ + *α*(*ϵ*_*orig*_ − *ϵ*_0_), where *α* = 0.7 was found by Benayad et al. to capture the phase separation behavior of FUS-LCD in the CG-Martini simulations. In the above formula, *ϵ*_*α*_ and *ϵ*_*orig*_ represent the scaled and the original LJ well depths, respectively, and *ϵ*_0_ = 2.0 kJ mol^−1^, which corresponds to the least attractive interaction in the Martini force field.^91^

The system was energy minimized and simulated for 1 *µ*s under *NpT* conditions at *T* = 300 K and *p* = 1 bar. Temperature and pressure were maintained using a weak coupling thermostat and barostat,^93^ respectively. All coarse-grained simulations were performed using the recommended settings,^94^ which included the use of a 20 fs time step to integrate the equations of motion.

From the obtained initial equilibrated system, batches of water molecules were removed in a step-wise manner to eventually obtain the desired protein concentrations (25, 60, 100, 145, 205, 260, 350, and 525 mg mL^−1^), see Figure 2 and Table 2. Before each next water removal step, the systems were further equilibrated under *NpT* conditions for 100 ns. The CG-Martini simulations enable fast equilibration of the protein configurations. However, the approximate and effective nature of the potential energy function and the reduced number of degrees of freedom (for example, the Martini water model does not have any rotational degrees of freedom) renders it difficult to obtain insights into the protein and solvent contributions to the thermodynamic driving forces. Therefore, the CG simulations were not used for any of the analyses presented in this work, but only to speed up the initial equilibration of the systems. To generate atomistic configurations for the subsequent all-atom MD simulations, the systems were back-mapped to the all-atom level using the *initram*.*sh* and *backward*.*py* scripts.^95^ The resulting atomistic proteins were solvated using the a99SB-disp water model, which was derived from the TIP4P-D model for usage together with the corresponding a99SB-disp protein force field,^96^ which was shown to be a good choice for FUS in a recent systematic comparison of nine force fields.^97^ The systems were neutralized by adding sodium ions, and additional 150 mM NaCl was added to mimic physiological ion concentration. The system details are summarized in Table 2. A separate cubic box of neat water, containing 177382 a99SB-disp water molecules and 150 mM NaCl, with box dimensions of ∼17.5 nm, was simulated to obtain the reference values for bulk water.

**Table 2:** Details of simulation systems studied in this work (CG = coarse grained; AA = all-atom).

| FUS conc.<br>(mg mL<sup>-1</sup>) | Box size<br>(nm) | Num. of water<br>beads/molecules |  | Num. of Na<sup>+</sup><br>ions |  | Num. of Cl<sup>-</sup><br>ions |  |
| --- | --- | --- | --- | --- | --- | --- | --- |
|  |  | CG | AA | CG | AA | CG | AA |
| 25 | 21.06 | 74366 | 304821 | 16 | 870 | 0 | 854 |
| 60 | 15.79 | 30366 | 125496 | 16 | 379 | 0 | 363 |
| 100 | 13.21 | 18366 | 71268 | 16 | 229 | 0 | 213 |
| 145 | 11.68 | 12366 | 47752 | 16 | 165 | 0 | 149 |
| 205 | 10.38 | 8366 | 32038 | 16 | 122 | 0 | 106 |
| 260 | 9.58 | 6366 | 24158 | 16 | 101 | 0 | 85 |
| 350 | 8.67 | 4366 | 16509 | 16 | 79 | 0 | 63 |
| 525 | 7.57 | 2366 | 9273 | 16 | 58 | 0 | 42 |

### MD simulation details

All systems were first energy minimized using steepest descent and then equilibrated with harmonic position restraints on the protein backbone for 10 ns (restraining force constants of 1000 kJ mol^−1^ nm^−2^) under *NpT* conditions at *T* = 300 K and *p* = 1 bar. This was followed by further *NpT* equilibration under the same conditions for 100 ns without any position restraints. Thereafter, the systems were equilibrated for 100 ns in the *NV T* ensemble, followed by the final production runs (see below). In these simulations, the velocity rescaling thermostat with a stochastic term^98^ and the Berendsen barostat^93^ were used to control temperature and pressure, respectively.

Two sets of production runs were carried out. For the vibrational density of states, 20 ps simulations with an output frequency of 4 fs were carried out. For the other calculations, 100 ns simulations with an output frequency of 1 ps were done. Every simulation was repeated 3 times, starting from different initial configurations. Statistical errors were calculated as standard deviations over these three trajectories. All atomistic MD simulations were done using the leap-frog integrator with a time step of 2 fs. Protein bonds and internal degrees of water molecules were constrained using the LINCS and SETTLE algorithms, respectively. Short-range electrostatic and Lennard-Jones interactions were calculated up to an interparticle distance cut-off of 1.0 nm. Long-range electrostatic interactions were treated with the particle-mesh Ewald technique^99^ with a grid spacing of 0.12 nm.

### Analyses

#### Interaction energy

The interaction energies were calculated by post-processing the simulation trajectories using the *mdrun -rerun* routine implemented in GROMACS, with Coulomb and van der Waals distance cut-offs increased to 3.5 nm. Such a large cut-off distance of the pairwise nonbonded interactions was used to minimize the neglect of the lattice part (long-range interactions) in the reruns.

#### Entropy of water

The entropy of water was computed using the the DoSPT implementation^100,101^ of the 2-phase-thermodynamics (2PT) method. ^77,102^ This method was specifically developed to calculate the molar entropies of liquids, and it was shown to yield accurate water entropies in different systems and under a variety of conditions. ^21,24,103–107^ In 2PT, the spectral density *I*(*ν*) or density of states (DoS) of a liquid is the Fourier transform of the velocity autocorrelation function (VACF), Eq. 18.

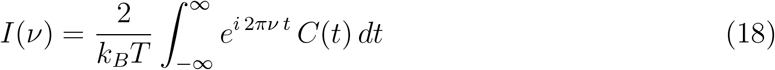

For rigid water molecules, translational (*C*_*tr*_(*t*)) and rotational (*C*_*rot*_(*t*)) VACFs can be treated separately,

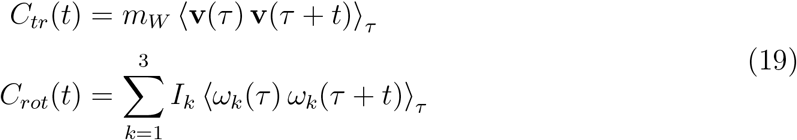

where *m*_*W*_ is the mass of a water molecule, v its center-of-mass velocity vector, *I*_*k*_ the moment of inertia for rotation around the *k*-th principal axis of the water molecule, *ω*_*k*_ the corresponding angular velocity, and ⟨…⟩_*τ*_ is an ensemble average over initial times *τ* . The 2PT method partitions the total DoS of the liquid into a solid-like (*I*^*s*^(*ν*)) and a gas-like (*I*^*g*^(*ν*)) contribution,

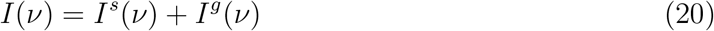

The solid-like DoS is treated using the harmonic oscillator (HO) model. The diffusive gas-like contribution is described using the Enskog hard sphere (HS) theory (for translation) and the rigid rotor (RR) model (for rotation). The analytically obtained translational and rotational contributions are added to arrive at the total entropy of water (Eq. 21),

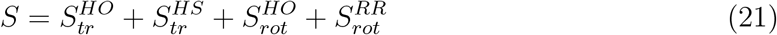

An advantage of the 2PT approach is that it yields translational and rotational entropy separately. For more detailed descriptions of the method, see the works of Lin et al. ^77,102^ and the recent review by Heyden.^24^

#### Protein Conformational Entropy

We estimated the conformational entropy of the proteins using a nearest neighbor (NN) approach,^108,109^ as implemented in the *PDB2ENTROPY* method introduced by Fogolari et al. that estimates conformational entropies of proteins from probability distributions of torsion angles relative to uniform distributions.^81^ The torsion angles are considered for atoms that are within a given cut-off distance (8 °A) from the central atom. Entropy was calculated based on the maximum information spanning tree (MIST) approach.^110^ To calculate the conformational entropy difference, ∆*S*_*conf*_, between the condensed and dilute phases, the simulation systems at 350 mg/mL and 25 mg/mL concentration were used. We averaged over all 8 FUS-LCD proteins in the simulation boxes and over the three independent repeats.

All the software and codes used in work are tabulated in Table S11 in the Supporting Information.

## Data Availability

Data supporting the findings of this manuscript are available from the corresponding authors upon request. All data plotted in Figures 4, 7, 8 are provided as Tables in Supporting Information, which also contains the initial and final configurations of the MD simulations, MD-parameter files, and force field topology files.

## Code Availability

The computer codes used to run and analyze the MD simulations are available in the public domain, see list in Table S11 in Supporting Information.

## Acknowledgements

We thank Matthias Heyden (Arizona State University) for useful discussions. This project received funding from the European Union’s Horizon 2020 research and innovation programme under the Marie Skłodowska-Curie grant agreement No 801459 - FP-RESOMUS and from the Deutsche Forschungsgemeinschaft (DFG) under Germany’s Excellence Strategy - EXC 2033 - 390677874 - RESOLV.

## Author Contributions

L.V.S. conceived the study and designed the project. S.M. carried out all MD simulations, analyzed and interpreted the data, and created the figures, with the help of L.V.S. Both authors wrote the manuscript.

## References

(1) Zimmerman, S. B.; Trach, S. O. Estimation of macromolecule concentrations and excluded volume effects for the cytoplasm of Escherichia coli. J. Mol. Biol. 1991, 222, 599–620.

(2) Wang, Y.; Sarkar, M.; Smith, A. E.; Krois, A. S.; Pielak, G. J. Macromolecular crowding and protein stability. J. Am. Chem. Soc. 2012, 134, 16614–16618.

(3) Gao, M.; Held, C.; Patra, S.; Arns, L.; Sadowski, G.; Winter, R. Crowders and cosolvents—major contributors to the cellular milieu and efficient means to counteract environmental stresses. ChemPhysChem 2017, 18, 2951–2972.

(4) Theillet, F.-X.; Binolfi, A.; Frembgen-Kesner, T.; Hingorani, K.; Sarkar, M.; Kyne, C.; Li, C.; Crowley, P. B.; Gierasch, L.; Pielak, G. J., et al. Physicochemical properties of cells and their effects on intrinsically disordered proteins (IDPs). Chem. Rev. 2014, p114, 6661–6714.

(5) Barbosa, A. D.; Savage, D. B.; Siniossoglou, S. Lipid droplet–organelle interactions: emerging roles in lipid metabolism. Curr. Opin. Cell Biol. 2015, p35, 91–97.

(6) Speer, S. L.; Zheng, W.; Jiang, X.; Chu, I.-T.; Guseman, A. J.; Liu, M.; Pielak, G. J.; Li, C. The intracellular environment affects protein–protein interactions. Proc. Natl. Acad. Sci. U.S.A. 2021, 118, e2019918118.

(7) Legrain, P.; Wojcik, J.; Gauthier, J.-M. Protein–protein interaction maps: a lead towards cellular functions. Trends Genet. 2001, p17, 346–352.

(8) Lasker, K.; Boeynaems, S.; Lam, V.; Scholl, D.; Stainton, E.; Briner, A.; Jacquemyn, M.; Daelemans, D.; Deniz, A.; Villa, E., et al. The material properties of a bacterial-derived biomolecular condensate tune biological function in natural and synthetic systems. Nat. Commun. 2022, 13, 5643.

(9) Sahni, N.; Yi, S.; Taipale, M.; Bass, J. I. F.; Coulombe-Huntington, J.; Yang, F.; Peng, J.; Weile, J.; Karras, G. I.; Wang, Y., et al. Widespread macromolecular interaction perturbations in human genetic disorders. Cell 2015, p161, 647–660.

(10) Grudzielanek, S.; Smirnovas, V.; Winter, R. Solvation-assisted pressure tuning of insulin fibrillation: from novel aggregation pathways to biotechnological applications. J. Mol. Biol. 2006, p356, 497–509.

(11) Chiti, F.; Dobson, C. M., et al. Protein misfolding, functional amyloid, and human disease. Annu. Rev. Biochem. 2006, p75, 333–366.

(12) Thirumalai, D.; Reddy, G.; Straub, J. E. Role of water in protein aggregation and amyloid polymorphism. Acc. Chem. Res. 2012, p45, 83–92.

(13) Bentmann, E.; Neumann, M.; Tahirovic, S.; Rodde, R.; Dormann, D.; Haass, C. Requirements for stress granule recruitment of fused in sarcoma (FUS) and TAR DNA-binding protein of 43 kDa (TDP-43). J. Biol. Chem. 2012, p287, 23079–23094.

(14) Ramaswami, M.; Taylor, J. P.; Parker, R. Altered ribostasis: RNA-protein granules in degenerative disorders. Cell 2013, 154, 727–736.

(15) Patel, A.; Lee, H. O.; Jawerth, L.; Maharana, S.; Jahnel, M.; Hein, M. Y.; Stoynov, S.; Mahamid, J.; Saha, S.; Franzmann, T. M., et al. A liquid-to-solid phase transition of the ALS protein FUS accelerated by disease mutation. Cell 2015, 162, 1066–1077.

(16) Hofweber, M.; Hutten, S.; Bourgeois, B.; Spreitzer, E.; Niedner-Boblenz, A.; Schifferer, M.; Ruepp, M.-D.; Simons, M.; Niessing, D.; Madl, T., et al. Phase separation of FUS is suppressed by its nuclear import receptor and arginine methylation. Cell 2018, p173, 706–719.

(17) Dignon, G. L.; Best, R. B.; Mittal, J. Biomolecular phase separation: From molecular driving forces to macroscopic properties. Annu. Rev. Phys. Chem. 2020, 71, 53–75.

(18) Bagchi, B. Water in biological and chemical processes: From structure and dynamics to function; Cambridge University Press, 2013.

(19) Bellissent-Funel, M.-C.; Hassanali, A.; Havenith, M.; Henchman, R.; Pohl, P.; Sterpone, F.; van der Spoel, D.; Xu, Y.; Garcia, A. E. Water determines the structure and dynamics of proteins. Chem. Rev. 2016, 116, 7673–7697.

(20) Mondal, S.; Bagchi, B. From structure and dynamics to biomolecular functions: The ubiquitous role of solvent in biology. Curr. Opin. Struct. Biol. 2022, 77, 102462.

(21) Mukherjee, S.; Schäfer, L. V. Spatially Resolved Hydration Thermodynamics in Biomolecular Systems. J. Phys. Chem. B 2022, 126, 3619–3631.

(22) Mukherjee, S.; Mondal, S.; Acharya, S.; Bagchi, B. Tug-of-War between Internal and External Frictions and Viscosity Dependence of Rate in Biological Reactions. Phys. Rev. Lett. 2022, p128, 108101.

(23) Ribeiro, S. S.; Samanta, N.; Ebbinghaus, S.; Marcos, J. C. The synergic effect of water and biomolecules in intracellular phase separation. Nat. Rev. Chem. 2019, 3, 552–561.

(24) Heyden, M. Disassembling solvation free energies into local contributions – Toward a microscopic understanding of solvation processes. WIREs Comput. Mol. Sci. 2019, p9, e1390.

(25) Alberti, S.; Gladfelter, A.; Mittag, T. Considerations and challenges in studying liquidliquid phase separation and biomolecular condensates. Cell 2019, p176, 419–434.

(26) Martin, E. W.; Holehouse, A. S.; Peran, I.; Farag, M.; Incicco, J. J.; Bremer, A.; Grace, C. R.; Soranno, A.; Pappu, R. V.; Mittag, T. Valence and patterning of aromatic residues determine the phase behavior of prion-like domains. Science 2020, p367, 694–699.

(27) Mittag, T.; Pappu, R. V. A conceptual framework for understanding phase separation and addressing open questions and challenges. Mol. Cell 2022, p82, 2201–2214.

(28) Pappu, R. V.; Cohen, S. R.; Dar, F.; Farag, M.; Kar, M. Phase Transitions of Associative Biomacromolecules. Chem. Rev. 2023, p123, 8945–8987.

(29) Brangwynne, C. P.; Eckmann, C. R.; Courson, D. S.; Rybarska, A.; Hoege, C.; Gharakhani, J.; Jülicher, F.; Hyman, A. A. Germline P granules are liquid droplets that localize by controlled dissolution/condensation. Science 2009, 324, 1729–1732.

(30) Banani, S. F.; Lee, H. O.; Hyman, A. A.; Rosen, M. K. Biomolecular condensates: organizers of cellular biochemistry. Nat. Rev. Mol. Cell Biol. 2017, 18, 285–298.

(31) Krainer, G.; Welsh, T. J.; Joseph, J. A.; Espinosa, J. R.; Wittmann, S.; de Csilléry, E.; Sridhar, A.; Toprakcioglu, Z.; Gudiškyte?, G.; Czekalska, M. A., et al. Reentrant liquid condensate phase of proteins is stabilized by hydrophobic and non-ionic interactions. Nat. Commun. 2021, 12, 1085.

(32) Banjade, S.; Rosen, M. K. Phase transitions of multivalent proteins can promote clustering of membrane receptors. eLife 2014, 3, e04123.

(33) Shin, Y.; Brangwynne, C. P. Liquid phase condensation in cell physiology and disease.\ Science 2017, 357, eaaf4382.

(34) Strom, A. R.; Emelyanov, A. V.; Mir, M.; Fyodorov, D. V.; Darzacq, X.; Karpen, G. H. Phase separation drives heterochromatin domain formation. Nature 2017, p547, 241–245.

(35) Wang, A.; Conicella, A. E.; Schmidt, H. B.; Martin, E. W.; Rhoads, S. N.; Reeb, A. N.; Nourse, A.; Ramirez Montero, D.; Ryan, V. H.; Rohatgi, R., et al. A single N-terminal phosphomimic disrupts TDP-43 polymerization, phase separation, and RNA splicing. EMBO J. 2018, 37, e97452.

(36) Woodruff, J. B. Assembly of mitotic structures through phase separation. J. Mol. Biol. 2018, p430, 4762–4772.

(37) Brangwynne, C. P.; Mitchison, T. J.; Hyman, A. A. Active liquid-like behavior of nucleoli determines their size and shape in Xenopus laevis oocytes. Proc. Natl. Acad. Sci. U.S.A. 2011, p108, 4334–4339.

(38) Li, Q.; Peng, X.; Li, Y.; Tang, W.; Zhu, J.; Huang, J.; Qi, Y.; Zhang, Z. LLPSDB: a database of proteins undergoing liquid–liquid phase separation in vitro. Nucleic Acids Res. 2020, p48, D320–D327.

(39) Wang, X.; Zhou, X.; Yan, Q.; Liao, S.; Tang, W.; Xu, P.; Gao, Y.; Li, Q.; Dou, Z.; Yang, W., et al. LLPSDB v2.0: an updated database of proteins undergoing liquid– liquid phase separation in vitro. Bioinformatics 2022, p38, 2010–2014.

(40) Burke, K. A.; Janke, A. M.; Rhine, C. L.; Fawzi, N. L. Residue-by-residue view of in vitro FUS granules that bind the C-terminal domain of RNA polymerase II. Mol. Cell 2015, 60, 231–241.

(41) Murakami, T.; Qamar, S.; Lin, J. Q.; Schierle, G. S. K.; Rees, E.; Miyashita, A.; Costa, A. R.; Dodd, R. B.; Chan, F. T.; Michel, C. H., et al. ALS/FTD mutation-induced phase transition of FUS liquid droplets and reversible hydrogels into irreversible hydrogels impairs RNP granule function. Neuron 2015, p88, 678–690.

(42) Kitahara, R.; Yamazaki, R.; Ide, F.; Li, S.; Shiramasa, Y.; Sasahara, N.; Yoshizawa, T. Pressure-Jump Kinetics of Liquid–Liquid Phase Separation: Comparison of Two Different Condensed Phases of the RNA-Binding Protein, Fused in Sarcoma. J. Am. Chem. Soc. 2021, p143, 19697–19702.

(43) Guo, W.; Naujock, M.; Fumagalli, L.; Vandoorne, T.; Baatsen, P.; Boon, R.; Ordovás, L.; Patel, A.; Welters, M.; Vanwelden, T., et al. HDAC6 inhibition reverses axonal transport defects in motor neurons derived from FUS-ALS patients. Nat. Commun. 2017, 8, 1–15.

(44) Kamelgarn, M.; Chen, J.; Kuang, L.; Jin, H.; Kasarskis, E. J.; Zhu, H. ALS mutations of FUS suppress protein translation and disrupt the regulation of nonsense-mediated decay. Proc. Natl. Acad. Sci. U.S.A. 2018, 115, E11904–E11913.

(45) Rulten, S. L.; Rotheray, A.; Green, R. L.; Grundy, G. J.; Moore, D. A. Q.; Gómez-Herreros, F.; Hafezparast, M.; Caldecott, K. W. PARP-1 dependent recruitment of the amyotrophic lateral sclerosis-associated protein FUS/TLS to sites of oxidative DNA damage. Nucleic Acids Res. 2013, 42, 307–314.

(46) Zhang, T. et al. FUS Regulates Activity of MicroRNA-Mediated Gene Silencing. Mol. Cell 2018, p69, 787–801.

(47) Yasuda, K.; Zhang, H.; Loiselle, D.; Haystead, T.; Macara, I. G.; Mili, S. The RNAbinding protein Fus directs translation of localized mRNAs in APC-RNP granules. J. Cell Biol. 2013, 203, 737–746.

(48) Yokosawa, K.; Kajimoto, S.; Shibata, D.; Kuroi, K.; Konno, T.; Nakabayashi, T. Concentration Quantification of the Low-Complexity Domain of Fused in Sarcoma inside a Single Droplet and Effects of Solution Parameters. J. Phys. Chem. Lett. 2022, 13, 5692–5697.

(49) Benayad, Z.; von Bülow, S.; Stelzl, L. S.; Hummer, G. Simulation of FUS protein condensates with an adapted coarse-grained model. J. Chem. Theory Comput. 2021, p17, 525–537.

(50) Avni, A.; Joshi, A.; Walimbe, A.; Pattanashetty, S. G.; Mukhopadhyay, S. Single-droplet surface-enhanced Raman scattering decodes the molecular determinants of liquid-liquid phase separation. Nat. Commun. 2022, 13, 4378.

(51) Nott, T. J.; Petsalaki, E.; Farber, P.; Jervis, D.; Fussner, E.; Plochowietz, A.; Craggs, T. D.; Bazett-Jones, D. P.; Pawson, T.; Forman-Kay, J. D., et al. Phase transition of a disordered nuage protein generates environmentally responsive membraneless organelles. Mol. Cell 2015, 57, 936–947.

(52) Lin, Y.; McCarty, J.; Rauch, J. N.; Delaney, K. T.; Kosik, K. S.; Fredrickson, G. H.; Shea, J.-E.; Han, S. Narrow equilibrium window for complex coacervation of tau and RNA under cellular conditions. eLife 2019, 8, e42571.

(53) Ahlers, J.; Adams, E. M.; Bader, V.; Pezzotti, S.; Winklhofer, K. F.; Tatzelt, J.; Havenith, M. The key role of solvent in condensation: mapping water in liquid-liquid phase-separated FUS. Biophys. J. 2021, 120, 1266–1275.

(54) Pezzotti, S.; König, B.; Ramos, S.; Schwaab, G.; Havenith, M. Liquid–Liquid Phase Separation? Ask the Water! J. Phys. Chem. Lett. 2023, 14, 1556–1563.

(55) Murthy, A. C.; Dignon, G. L.; Kan, Y.; Zerze, G. H.; Parekh, S. H.; Mittal, J.; Fawzi, N. L. Molecular interactions underlying liquidliquid phase separation of the FUS low-complexity domain. Nat. Struct. Mol. Biol. 2019, p26, 637–648.

(56) Shea, J.-E.; Best, R. B.; Mittal, J. Physics-based computational and theoretical approaches to intrinsically disordered proteins. Curr. Opin. Struct. Biol. 2021, 67, 219–225.

(57) Zheng, W.; Dignon, G. L.; Jovic, N.; Xu, X.; Regy, R. M.; Fawzi, N. L.; Kim, Y. C.; Best, R. B.; Mittal, J. Molecular Details of Protein Condensates Probed by Microsecond Long Atomistic Simulations. J. Phys. Chem. B 2020, 124, 11671–11679.

(58) Paloni, M.; Bailly, R.; Ciandrini, L.; Barducci, A. Unraveling Molecular Interactions in Liquid–Liquid Phase Separation of Disordered Proteins by Atomistic Simulations. J. Phys. Chem. B 2020, 124, 9009–9016.

(59) Guseva, S.; Schnapka, V.; Adamski, W.; Maurin, D.; Ruigrok, R. W. H.; Salvi, N.; Blackledge, M. Liquid–Liquid Phase Separation Modifies the Dynamic Properties of Intrinsically Disordered Proteins. J. Am. Chem. Soc. 2023, p145, 10548–10563.

(60) Galvanetto, N.; Ivanović, M. T.; Chowdhury, A.; Sottini, A.; Nüesch, M. F.; Nettels, D.; Best, R. B.; Schuler, B. Extreme dynamics in a biomolecular condensate. Nature 2023, p619, 876–883.

(61) Harmon, T. S.; Holehouse, A. S.; Rosen, M. K.; Pappu, R. V. Intrinsically disordered linkers determine the interplay between phase separation and gelation in multivalent proteins. eLife 2017, p6, e30294.

(62) Dignon, G. L.; Zheng, W.; Kim, Y. C.; Best, R. B.; Mittal, J. Sequence determinants of protein phase behavior from a coarse-grained model. PLOS Comput. Biol. 2018, 14, 1–23.

(63) Dignon, G. L.; Zheng, W.; Best, R. B.; Kim, Y. C.; Mittal, J. Relation between singlemolecule properties and phase behavior of intrinsically disordered proteins. Proc. Natl. Acad. Sci. U.S.A. 2018, p115, 9929–9934.

(64) Statt, A.; Casademunt, H.; Brangwynne, C. P.; Panagiotopoulos, A. Z. Model for disordered proteins with strongly sequence-dependent liquid phase behavior. J. Chem. Phys. 2020, 152 .

(65) Regy, R. M.; Zheng, W.; Mittal, J. Using a sequence-specific coarse-grained model for studying protein liquid–liquid phase separation. Methods Enzymol. 2021, 646, 1–17.

(66) McCall, P. M.; Kim, K.; Fritsch, A. W.; Iglesias-Artola, J.; Jawerth, L. M.; Wang, J.; Ruer, M.; Peychl, J.; Poznyakovskiy, A.; Guck, J.; Alberti, S.; Hyman, A. A.; Brugués, J. Quantitative phase microscopy enables precise and efficient determination of biomolecular condensate composition. bioRxiv 2020,

(67) Murakami, K.; Kajimoto, S.; Shibata, D.; Kuroi, K.; Fujii, F.; Nakabayashi, T. Observation of liquid–liquid phase separation of ataxin-3 and quantitative evaluation of its concentration in a single droplet using Raman microscopy. Chem. Sci. 2021, 12, 7411–7418.

(68) Chau, P.-L.; Hardwick, A. A new order parameter for tetrahedral configurations. Mol. Phys. 1998, 93, 511–518.

(69) Errington, J. R.; Debenedetti, P. G. Relationship between structural order and the anomalies of liquid water. Nature 2001, 409, 318–321.

(70) Duboué-Dijon, E.; Laage, D. Characterization of the local structure in liquid water by various order parameters. J. Phys. Chem. B 2015, p119, 8406–8418.

(71) Wang, H.; Kelley, F. M.; Milovanovic, D.; Schuster, B. S.; Shi, Z. Surface tension and viscosity of protein condensates quantified by micropipette aspiration. Biophys. Rep. 2021, p1, 100011.

(72) Wang, B.; Zhang, L.; Dai, T.; Qin, Z.; Lu, H.; Zhang, L.; Zhou, F. Liquid–liquid phase separation in human health and diseases. Sig. Transduct. Target Ther. 2021, p6, 290.

(73) Ben-Naim, A. Hydrophobic interaction and structural changes in the solvent. Biopolymers. 1975, p14, 1337–1355.

(74) Yu, H.-A.; Karplus, M. A thermodynamic analysis of solvation. J. Chem. Phys. 1988, 89, 2366–2379.

(75) Ben-Amotz, D. Water-mediated hydrophobic interactions. Annu. Rev. Phys. Chem. 2016, 67, 617–638.

(76) Heinz, L. P.; Grubmüller, H. Why solvent response contributions to solvation free energies are compatible with Ben-Naim’s theorem. arXiv 2023, 2306.09392.

(77) Lin, S.-T.; Maiti, P. K.; Goddard III, W. A. Two-phase thermodynamic model for efficient and accurate absolute entropy of water from molecular dynamics simulations. J. Phys. Chem. B 2010, p114, 8191–8198.

(78) Zwanzig, R. W. High-temperature equation of state by a perturbation method. I. Nonpolar gases. J. Chem. Phys. 1954, 22, 1420–1426.

(79) Shirts, M. R.; Pande, V. S. Solvation free energies of amino acid side chain analogs for common molecular mechanics water models. J. Chem. Phys. 2005, 122, 134508.

(80) Molliex, A.; Temirov, J.; Lee, J.; Coughlin, M.; Kanagaraj, A. P.; Kim, H. J.; Mittag, T.; Taylor, J. P. Phase separation by low complexity domains promotes stress granule assembly and drives pathological fibrillization. Cell 2015, 163, 123–133.

(81) Fogolari, F.; Maloku, O.; Dongmo Foumthuim, C. J.; Corazza, A.; Esposito, G. PDB2ENTROPY and PDB2TRENT: Conformational and translational–rotational entropy from molecular ensembles. J. Chem. Inf. Model. 2018, 58, 1319–1324.

(82) Genheden, S.; Ryde, U. Will Molecular Dynamics Simulations of Proteins Ever Reach Equilibrium? Phys. Chem. Chem. Phys. 2012, 14, 8662–8677.

(83) Hoffmann, F.; Mulder, F. A. A.; Schäfer, L. V. How Much Entropy Is Contained in NMR Relaxation Parameters? J. Phys. Chem. B 2022, 126, 54–68.

(84) Boeynaems, S.; Holehouse, A. S.; Weinhardt, V.; Kovacs, D.; Lindt, J. V.; Larabell, C.; Bosch, L. V. D.; Das, R.; Tompa, P. S.; Pappu, R. V.; Gitler, A. D. Spontaneous driving forces give rise to protein-RNA condensates with coexisting phases and complex material properties. Proc. Natl. Acad. Sci. U.S.A. 2019, p116, 7889–7898.

(85) Erkamp, N. A.; Sneideris, T.; Ausserwöger, H.; Qian, D.; Qamar, S.; Nixon-Abell, J.; St George-Hyslop, P.; Schmit, J. D.; Weitz, D. A.; Knowles, T. P. J. Spatially nonuniform condensates emerge from dynamically arrested phase separation. Nat. Commun. 2023, 14, 684.

(86) Berendsen, H. J.; van der Spoel, D.; van Drunen, R. GROMACS: A message-passing parallel molecular dynamics implementation. Comput. Phys. Commun. 1995, p91, 43–56.

(87) Abraham, M. J.; Murtola, T.; Schulz, R.; Páll, S.; Smith, J. C.; Hess, B.; Lindahl, E. GROMACS: High performance molecular simulations through multi-level parallelism from laptops to supercomputers. SoftwareX 2015, p1, 19–25.

(88) Jumper, J.; Evans, R.; Pritzel, A.; Green, T.; Figurnov, M.; Ronneberger, O.; Tunyasuvunakool, K.; Bates, R.; Žídek, A.; Potapenko, A., et al. Highly accurate protein structure prediction with AlphaFold. Nature 2021, 596, 583–589.

(89) Varadi, M.; Anyango, S.; Deshpande, M.; Nair, S.; Natassia, C.; Yordanova, G.; Yuan, D.; Stroe, O.; Wood, G.; Laydon, A., et al. AlphaFold Protein Structure Database: Massively expanding the structural coverage of protein-sequence space with high-accuracy models. Nucleic Acids Res. 2022, 50, D439–D444.

(90) Monticelli, L.; Kandasamy, S. K.; Periole, X.; Larson, R. G.; Tieleman, D. P.; Marrink, S. J. The MARTINI coarse-grained force field: extension to proteins. J. Chem. Theory Comput. 2008, 4, 819–834.

(91) de Jong, D. H.; Singh, G.; Bennett, W. F. D.; Arnarez, C.; Wassenaar, T. A.; Schäfer, L. V.; Periole, X.; Tieleman, D. P.; Marrink, S. J. Improved parameters for the Martini coarse-grained protein force field. J. Chem. Theory Comput. 2013, 9, 687–697.

(92) Stark, A. C.; Andrews, C. T.; Elcock, A. H. Toward optimized potential functions for protein–protein interactions in aqueous solutions: osmotic second virial coefficient calculations using the martini coarse-grained force field. J. Chem. Theory Comput. 2013, 9, 4176–4185.

(93) Berendsen, H. J. C.; Postma, J. P. M.; van Gunsteren, W. F.; DiNola, A.; Haak, J. R. Molecular dynamics with coupling to an external bath. J. Chem. Phys. 1984, p81, 3684–3690.

(94) de Jong, D. H.; Baoukina, S.; Ingólfsson, H. I.; Marrink, S. J. Martini straight: Boosting performance using a shorter cutoff and GPUs. Comput. Phys. Commun. 2016, p199, 1–7.

(95) Wassenaar, T. A.; Pluhackova, K.; Böckmann, R. A.; Marrink, S. J.; Tieleman, D. P. Going backward: a flexible geometric approach to reverse transformation from coarse grained to atomistic models. J. Chem. Theory Comput. 2014, 10, 676–690.

(96) Robustelli, P.; Piana, S.; Shaw, D. E. Developing a molecular dynamics force field for both folded and disordered protein states. Proc. Natl. Acad. Sci. U.S.A. 2018, 115, E4758–E4766.

(97) Sarthak, K.; Winogradoff, D.; Ge, Y.; Myong, S.; Aksimentiev, A. Benchmarking Molecular Dynamics Force Fields for All-Atom Simulations of Biological Condensates. J. Chem. Theory Comput. 2023, 12, 3721–3740.

(98) Bussi, G.; Donadio, D.; Parrinello, M. Canonical sampling through velocity rescaling. J. Chem. Phys. 2007, 126, 014101.

(99) Darden, T.; York, D.; Pedersen, L. Particle mesh Ewald: An Nlog(N) method for Ewald sums in large systems. J. Chem. Phys. 1993, 98, 10089–10092.

(100) Caro, M. A.; Laurila, T.; Lopez-Acevedo, O. Accurate schemes for calculation of thermodynamic properties of liquid mixtures from molecular dynamics simulations. J. Chem. Phys. 2016, 145, 244504.

(101) Caro, M. A.; Lopez-Acevedo, O.; Laurila, T. Redox potentials from ab initio molecular dynamics and explicit entropy calculations: Application to transition metals in aqueous solution. J. Chem. Theory Comput. 2017, 13, 3432–3441.

(102) Lin, S.-T.; Blanco, M.; Goddard III, W. A. The two-phase model for calculating thermodynamic properties of liquids from molecular dynamics: Validation for the phase diagram of Lennard-Jones fluids. J. Chem. Phys. 2003, 119, 11792–11805.

(103) Persson, R. A.; Pattni, V.; Singh, A.; Kast, S. M.; Heyden, M. Signatures of solvation thermodynamics in spectra of intermolecular vibrations. J. Chem. Theory Comput. 2017, p13, 4467–4481.

(104) Fisette, O.; Päslack, C.; Barnes, R.; Isas, J. M.; Langen, R.; Heyden, M.; Han, S.; Schäfer, L. V. Hydration dynamics of a peripheral membrane protein. J. Am. Chem. Soc. 2016, p138, 11526–11535.

(105) Mukherjee, S.; Bagchi, B. Theoretical analyses of pressure induced glass transition in water: Signatures of surprising diffusion-entropy scaling across the transition. Mol. Phys. 2021, e1930222.

(106) Fajardo, T. N.; Heyden, M. Dissecting the Conformational Free Energy of a Small Peptide in Solution. J. Phys. Chem. B 2021, p125, 4634–4644.

(107) Päslack, C.; Das, C. K.; Schlitter, J.; Schäfer, L. V. Spectrally resolved estimation of water entropy in the active site of human carbonic anhydrase II. J. Chem. Theory Comput. 2021, p17, 5409–5418.

(108) Wang, L.; Abel, R.; Friesner, R. A.; Berne, B. J. Thermodynamic properties of liquid water: an application of a nonparametric approach to computing the entropy of a neat fluid. J. Chem. Theory Comput. 2009, 5, 1462–1473.

(109) Huggins, D. J. Estimating translational and orientational entropies using the k-nearest neighbors algorithm. J. Chem. Theory Comput. 2014, p10, 3617–3625.

(110) King, B. M.; Tidor, B. MIST: Maximum Information Spanning Trees for dimension reduction of biological data sets. Bioinformatics 2009, 25, 1165–1172.

