## Supporting Information for "Thermodynamic Forces from Protein and Water Govern Condensate Formation of an Intrinsically Disordered Protein Domain"

### Linear fit of entropy data

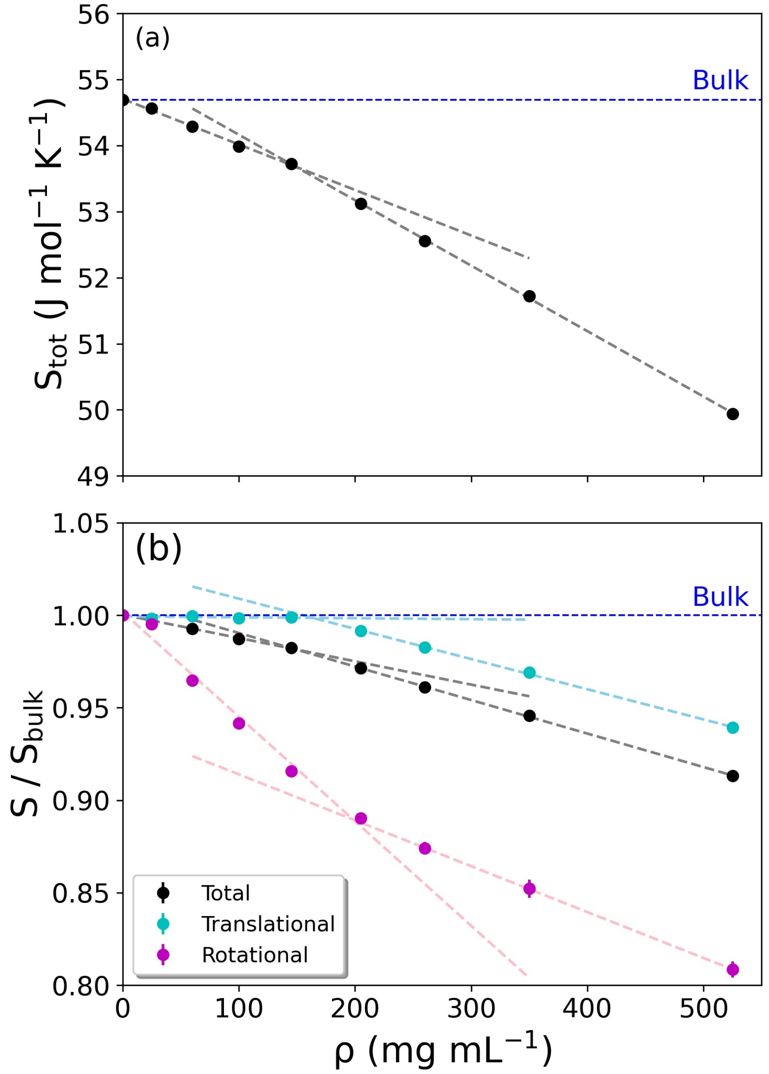


Figure S1: Entropy of water as a function of protein concentrations. Linear fits to the data are shown as dahsed lines. (a) Total entropy of water, (b) Translational, rotational and total entropy of water, each normalized with respect to bulk water.

### Distributions of radii of gyration

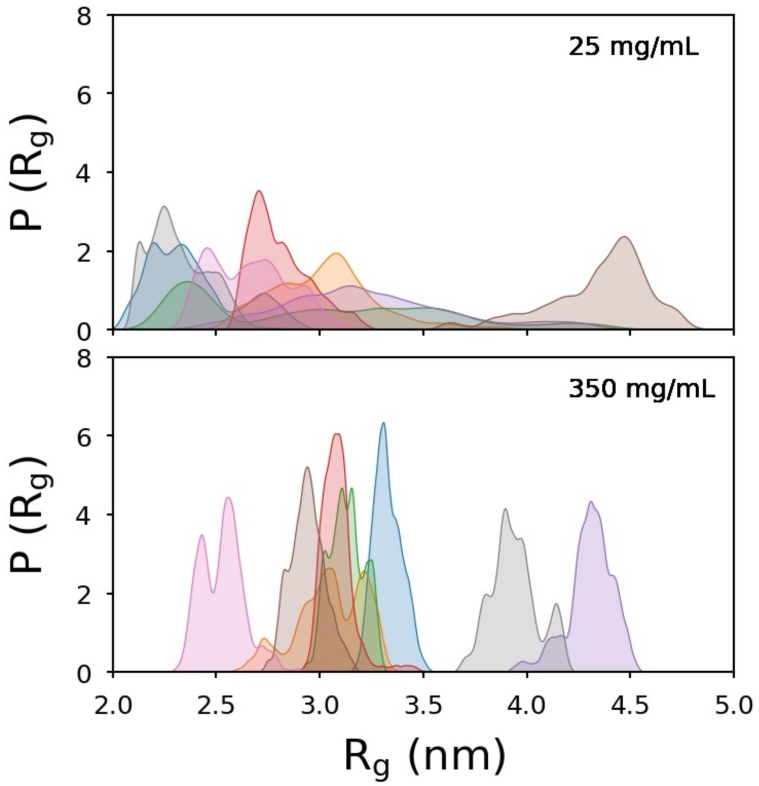


Figure S2: Distributions of the radii of gyration of the 8 FUS-LCD proteins present in the simulation boxes of the dilute (25 mg $mL^{-1}$) and the dense (350 mg $mL^{-1}$) systems. The distributions shown are based on the three 100 ns repeats, plus the data from the two trajectories that were extended to 400 ns.

### Water Entropy from Thermodynamic Integration/Free Energy Perturbation

From the total number of 177382 water molecules in the simulated cubic box of bulk (neat) a99sb-disp water, a single water molecule was randomly selected and thermodynamic integration was performed. This procedure, including the random selection of the perturbed water molecule, was repeated 100 times. To decouple the selected water molecule from the environment, initially the Coulombic interactions were turned off by changing the coupling parameter $\lambda$ from 0 to 1 in discrete steps of 0.1. This was followed by changing the van der Waals interactions in $\lambda$-steps of 0.05. A soft-core Lennard Jones potential was used with a scaling factor of 0.5. The equations of motion were integrated using Langevin dynamics with time-steps of 2 fs. The negative value of the obtained free energy of vaporization, averaged over the 100 repeats, gives the solvation free energy $\Delta G_{solv}$.

From thermodynamic integration, we obtained the value of $\Delta G_{solv}$ = -23.45 kJ mol$^{-1}$ for neat (bulk) a99SB-disp water at 300 K and 1 bar. The solvation free energy includes an analytical correction because of the difference in the reference states of the solvated and isolated (in vacuum) water molecules. The standard solvation free energy (and the standard enthalpy of vaporization etc.) describes the transfer of the solute (here, a water molecule) from a thermodynamic standard state of the gas (ideal gas at 1 bar) to the thermodynamic standard state in solution (ideal solution at 1 mol L$^{-1}$). This includes a change in concentration from 1 bar (that is, 1 mol per 24.94 L at 300 K) to 1 mol L$^{-1}$, which is not included in the TI calculation. The correction term is $w = -\int_{V_1}^{V_2} p\,dV = -RT\int_{V_1}^{V_2} \frac{1}{V} dV = -RT\ln(V_2/V_1) =$ 8.023 kJ mol$^{-1}$ and the corrected solvation free energy is $\Delta G_{solv}$ = -31.47 kJ mol$^{-1}$. Hence, the vaporization free energy $\Delta G_{vap}$= 31.47 kJ mol$^{-1}$.

The vaporization enthalpy is given by $\Delta H_{vap} = -E_{pot} + RT$, where $E_{pot}$ is the average interaction energy of a water molecule with its surrounding in bulk water. The obtained $E_{pot}$ from the simulation at a99SB-disp water is -52.53 kJ mol$^{-1}$. Since the water model used has fixed point charges, this potential energy needs to be corrected for the self-polarization, $\Delta E_{pol} = (\mu - \mu_0)^2)/2\alpha$. The magnitude of the dipole moment of the a99SB-disp water model is $\mu$ = 2.44 D and the dipole moment of a water in vacuum is $\mu_0$ = 1.85 D. With 1 D = $3.33 \times 10^{-30}$ C·m and the polarizability of water of $\alpha = 1.608 \times 10^{-40}$ F·m, one obtains $\Delta E_{pol}$ = 7.23 kJ mol$^{-1}$. Hence the corrected energy per water molecule is -45.30 kJ mol$^{-1}$ and with RT = 2.48 kJ mol$^{-1}$ one obtains $\Delta H_{vap}$ = 47.78 kJ mol$^{-1}$.

The entropy of water is thus obtained as $S = (\Delta H_{vap} - \Delta G_{vap})/T$. The entropy at 300 K is 54.4 J mol$^{-1}$ K$^{-1}$, in excellent agreement with the 2PT entropy of 54.7 J mol$^{-1}$ K$^{-1}$.

### Tables for changes in water population and thermodynamic quantities

This section tabulates the data plotted in Figures 4, 7, and 8 in the main text.

Table S1: Change in number of water molecules per protein.

| $\rho$ ($mg\ mL^{-1}$) | $N_W^{rele}$ | $N_W^{reta}$ |
|---|---|---|
| 25 | 36039 | 2064 |
| 60 | 13623 | 2064 |
| 100 | 6845 | 2064 |
| 145 | 3905 | 2064 |
| 170 | 2934 | 2064 |
| 205 | 1941 | 2064 |
| 260 | 956 | 2064 |
| 350 | 0 | 2064 |

Table S2: Change in total solvation entropy.

| $\rho$ ($mg\ mL^{-1}$) | $-T\Delta S_{solv}^{rele}$ ($kJ\ mol^{-1}$) | $-T\Delta S_{solv}^{reta}$ ($kJ\ mol^{-1}$) | $-T\Delta S_{solv}^{tot}$ ($kJ\ mol^{-1}$) |
|---|---|---|---|
| 25 | -1749.7 | 1738.2 | -11.5 |
| 60 | -1713.5 | 1578.9 | -134.6 |
| 100 | -1443.6 | 1403.2 | -40.4 |
| 145 | -1135.4 | 1238.5 | 103.1 |
| 205 | -913.2 | 867.6 | -45.6 |
| 260 | -611.9 | 517.7 | -94.3 |
| 350 | 0.0 | 0.0 | 0.0 |

Table S3: Change in protein-water entropy.

| $\rho$ ($mg\ mL^{-1}$) | $-T\Delta S_{PW}^{rele}$ ($kJ\ mol^{-1}$) | $-T\Delta S_{PW}^{reta}$ ($kJ\ mol^{-1}$) | $-T\Delta S_{PW}^{tot}$ ($kJ\ mol^{-1}$) |
|---|---|---|---|
| 25 | -13789.5 | 12872.5 | -917.1 |
| 60 | -12452.5 | 11775.8 | -676.7 |
| 100 | -10919.2 | 10370.1 | -549.1 |
| 145 | -9144.8 | 8829.9 | -314.9 |
| 205 | -6817.2 | 6414.6 | -402.6 |
| 260 | -4477.5 | 3998.1 | -479.4 |
| 350 | 0.0 | 0.0 | 0.0 |

Table S4: Change in water-water energy.

| $\rho$ $(mg\ mL^{-1})$ | $\Delta E_{WW}^{rele}$ $(kJ\ mol^{-1})$ | $\Delta E_{WW}^{reta}$ $(kJ\ mol^{-1})$ | $\Delta E_{WW}^{tot}$ $(kJ\ mol^{-1})$ |
|---|---:|---:|---:|
| 25 | -12039.8 | 11134.3 | -905.6 |
| 60 | -10739.0 | 10196.9 | -542.1 |
| 100 | -9475.6 | 8966.9 | -508.7 |
| 145 | -8009.4 | 7591.4 | -418.0 |
| 205 | -5904.0 | 5547.1 | -356.9 |
| 260 | -3865.6 | 3480.5 | -385.1 |
| 350 | 0.0 | 0.0 | 0.0 |

Table S5: Change in protein-water energy.

| $\rho$ $(mg\ mL^{-1})$ | $\Delta E_{PW}^{rele}$ $(kJ\ mol^{-1})$ | $\Delta E_{PW}^{reta}$ $(kJ\ mol^{-1})$ | $\Delta E_{PW}^{tot}$ $(kJ\ mol^{-1})$ |
|---|---:|---:|---:|
| 25 | 2791.4 | -2611.9 | 179.5 |
| 60 | 2475.1 | -2396.8 | 78.3 |
| 100 | 2189.7 | -2111.5 | 78.2 |
| 145 | 1846.3 | -1796.1 | 50.1 |
| 205 | 1371.8 | -1313.4 | 58.4 |
| 260 | 903.6 | -821.4 | 82.2 |
| 350 | 0.0 | 0.0 | 0.0 |

Table S6: Change in total solvation enthalpy.

| $\rho$ $(mg\ mL^{-1})$ | $\Delta H_{solv}^{rele}$ $(kJ\ mol^{-1})$ | $\Delta H_{solv}^{reta}$ $(kJ\ mol^{-1})$ | $\Delta H_{solv}^{tot}$ $(kJ\ mol^{-1})$ |
|---|---:|---:|---:|
| 25 | -9248.4 | 8522.4 | -726.0 |
| 60 | -8263.9 | 7800.2 | -463.7 |
| 100 | -7285.8 | 6855.4 | -430.5 |
| 145 | -6163.2 | 5795.3 | -367.9 |
| 205 | -4532.2 | 4233.7 | -298.5 |
| 260 | -2962.0 | 2659.0 | -302.9 |
| 350 | 0.0 | 0.0 | 0.0 |

Table S7: Change in solvation free energy.

| $\rho$ ($mg\ mL^{-1}$) | $\Delta G_{solv}^{rele}$ ($kJ\ mol^{-1}$) | $\Delta G_{solv}^{reta}$ ($kJ\ mol^{-1}$) | $\Delta G_{solv}^{tot}$ ($kJ\ mol^{-1}$) |
|---|---|---|---|
| 25 | -10998.1 | 10260.6 | -737.5 |
| 60 | -9977.4 | 9379.0 | -598.4 |
| 100 | -8729.4 | 8258.6 | -470.9 |
| 145 | -7298.5 | 7033.8 | -264.8 |
| 205 | -5445.4 | 5101.3 | -344.2 |
| 260 | -3573.9 | 3176.7 | -397.2 |
| 350 | 0.0 | 0.0 | 0.0 |

Table S8: Change in protein-protein energy.

| $\rho$ ($mg\ mL^{-1}$) | $\Delta E_{PP}$ ($kJ\ mol^{-1}$) |
|---|---|
| 25 | -815.5 |
| 60 | -430.4 |
| 100 | -358.0 |
| 145 | -175.9 |
| 205 | -233.0 |
| 260 | -307.4 |
| 350 | 0.0 |

Table S9: Change in protein conformational entropy

| $\rho$ ($mg\ mL^{-1}$) | $-T\Delta S_P$ ($kJ\ mol^{-1}$) |
|---|---|
| 25 | 165.3 |
| 60 | 117.5 |
| 100 | 97.8 |
| 145 | 88.8 |
| 205 | 115.4 |
| 260 | 97.8 |
| 350 | 0.0 |

Table S10: Change in total free energy

| $\rho$ ($mg\ mL^{-1}$) | $\Delta G$ ($kJ\ mol^{-1}$) |
|---|---|
| 25 | -1387.8 |
| 60 | -911.3 |
| 100 | -731.1 |
| 145 | -351.9 |
| 205 | -461.8 |
| 260 | -606.8 |
| 350 | 0.0 |

### Water population and thermodynamic changes with a FUS-LCD concentration of 525 mg mL$^{-1}$ in the condensate

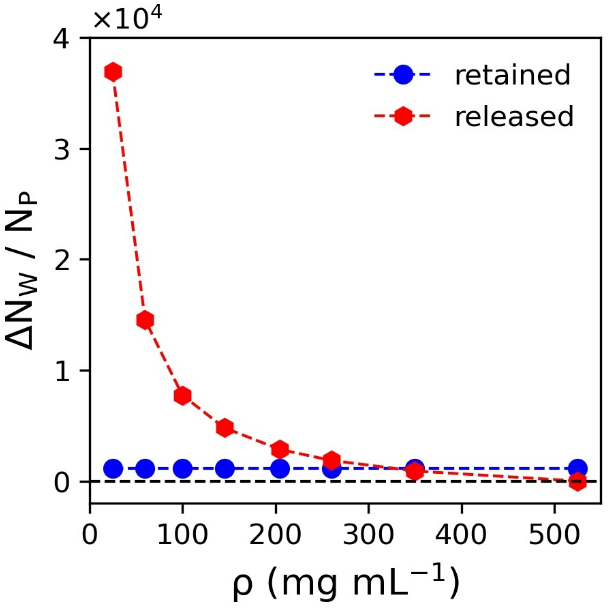


Figure S3: Number of released and retained water molecules per protein molecule as a function of protein concentration ($\rho$). The number of released molecules decreases (red line) as $\rho$ approaches the condensate concentration of 525 mg mL$^{-1}$, where every protein is solvated by 1159 water molecules, which are referred to as the retained waters (blue line).

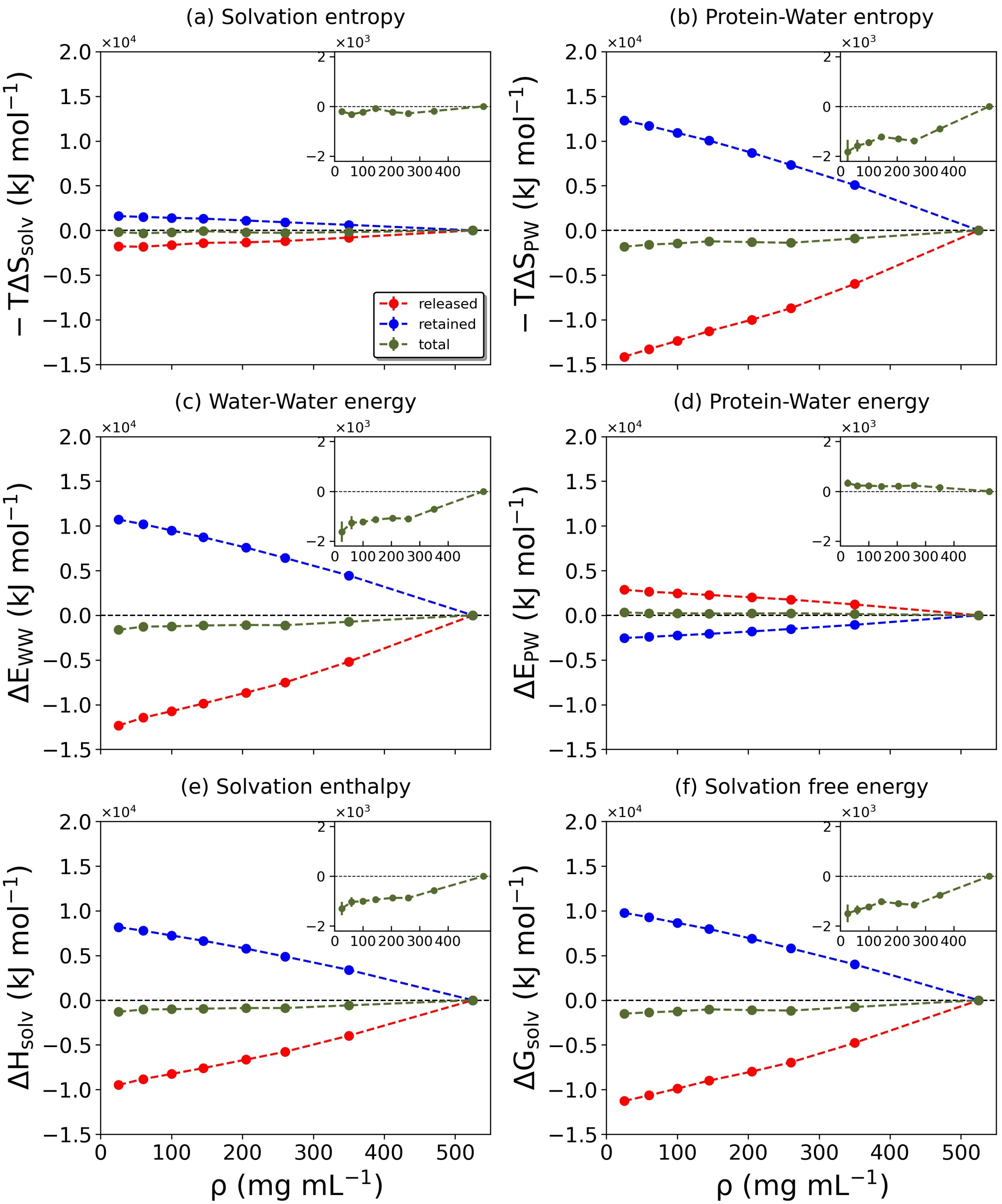


Figure S4: Changes of the solvation-related thermodynamic quantities. The quantities plotted in (a)-(f) are indicated at the top of each panel. The dashed red, blue, green lines denote the released, retained, and total water contributions, respectively. Data are presented as mean values ± standard deviation (SD) over three repeat simulations.

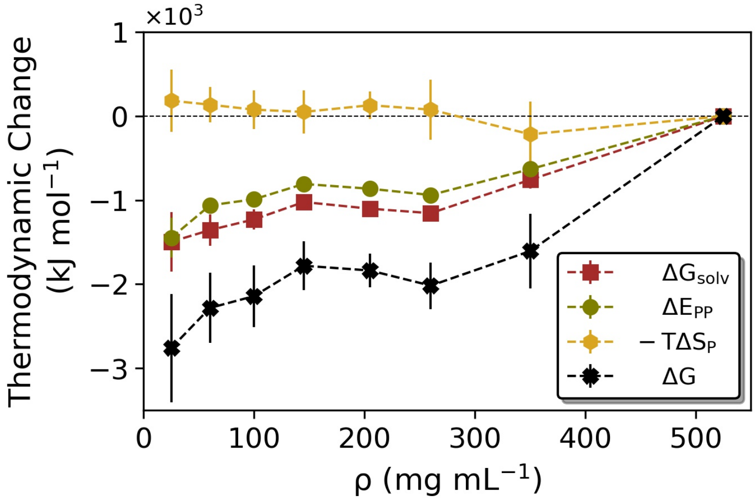


Figure S5: Thermodynamic contribution from changing total solvation free energy ($\Delta G_{solv}$, brown) protein-protein interaction energy ($\Delta E_{PP}$, green), and protein conformational entropy ($\Delta S_P$, orange) are plotted together with the resulting total free energy change ($\Delta G = \Delta G_{solv} + \Delta E_{PP} - T\Delta S_P$, black).

### Software/Codes used in this work

Table S11: Software/Codes used in this work, their functions and sources.

| Name of the code | Function | Source |
|---|---|---|
| DoSPT | Water entropy | www.dospt.org |
| pdb2entropy | Protein conformational entropy | https://github.com/ federico-fogolari/pdb2entropy |
| gmx mdrun -rerun | Interaction energies | |
| gmx energy | Interaction energies | |
| gmx hbond | number of hydrogen bonds | https://www.gromacs.org/ |
| gmx hydorder | tetrahedral order parameter | |
| gmx gyrate | radius of gyration | |
